# A c-opsin functions in a ciliary-marginal zone-like stem cell region of an invertebrate camera-type eye

**DOI:** 10.1101/2024.08.19.608633

**Authors:** Nadja Milivojev, Camila L. Velastegui Gamboa, Gabriele Andreatta, Florian Raible, Kristin Tessmar-Raible

**Author notes:** University of Padua, Department of Comparative Biomedicine and Food Science, Viale delĺUniversitá, 16, 35020 Legnaro, Italy (Gabriele Andreatta).

## Abstract

Camera-type eyes in vertebrates and invertebrates are striking examples of parallel evolution of a complex structure. Comparisons between such structures can help to deduce their organizational principles. We analyzed the camera-type adult eyes of the bristleworm *Platynereis dumerilii*. Employing single-cell RNA sequencing, we identified neurogenic cells in the worms’ adult brains. Among those are distinct neural stem cells in its adult eye, adjacent to the glass body/lens, that produce cells in radial lines, reminiscent of stem cells in the vertebrate eye ciliary marginal zone. A subset of these proliferating cells expresses the photoreceptor gene *c-opsin1*. *c-opsin1* knock-out reduces eye cell proliferation and influences differentiation. During reproductive maturation, proliferation in eye and brain sharply declines, while cells upregulate molecular characteristics of mammalian adult neural stem cell quiescence. Our data reveal new insights into nervous system functional development and evolution.

## Main Text

The characterization of adult neuronal stem cells (aNSCs) has advanced our insight into the neurogenic plasticity of vertebrate brains (*1*). In birds, the seasonal change from non-reproductive to reproductive stages is accompanied by changes in brain physiology driven by cell proliferation (*2*, *3*). In representatives of fishes or amphibians, aNSCs support life-long brain growth (*1*). Likewise, sensory organs such as the eye of these groups also contain a dedicated stem cell zone, termed the ciliary marginal zone (CMZ), from which new neurons and pigment cells differentiate (*4–6*). In amphibians and teleost fishes, retinal stem cells (RSCs) exhibit lasting growth, which may only decline with aging (*7*). By contrast, adult roles for RSCs are limited in birds, and lost in postnatal stages of mammals (*6*, *8*). In sharks, retinal neurogenesis is downregulated upon sexual maturation, possibly reflecting an ancestral regulation in jawed vertebrates (*9*). In the mammalian brain, aNSCs are not only confined to specific niches, but the ability of aNSCs to enter into neurogenic states is further limited by quiescence (*10*). Active *versus* quiescent neural stem cell states are characterized by specific molecular signatures of receptor signaling activity, transcription and translation (*11*).

Whereas this research attests to the importance of adult neuronal stem cells in the regulated growth of sensory organs and defined brain regions in vertebrates, there are only scarce comparative data for similar neurogenic processes in invertebrate model systems. In several invertebrate groups, enriched environmental stimuli can impact on neurogenesis in related processing centers: in red flour beetles, but not the fruitfly, the adult mushroom bodies retain some neurogenic potential, which depends on the olfactory environment (*12*). Similarly, environmental changes impact on adult neurogenesis in the olfactory lobes of the shore crab (*13*) as well as the octopus (*14*). Regulated changes in neurogenesis along an organism’s life history as they occur in vertebrates, however, are less explored in invertebrates. In ants, cast-specific hormone signatures lead to significant changes in the adult brain, mainly thought to occur via molecular regulation in differentiated neurons, not by changes in neurogenesis (*15*, *16*).

The marine bristleworm *Platynereis dumerilii* has emerged as a valuable model for cross-comparisons of the annelid nervous system with major bilaterian groups. Such comparisons have helped to clarify the evolutionary relationship of specific neuronal cell types. For instance, *Platynereis* larvae possess two molecularly distinct groups of photoreceptors that are demarcated by the expression of canonical rhabdomeric (*r-opsin*) and ciliary (*c-opsin*) opsin genes, respectively, the latter of which had long been assumed to be a vertebrate innovation (*17*).

Functional experiments imply *r-opsin1* – expressed in the adult eyes of the worms – in high-sensitivity light detection (*18*). By contrast, *c-opsin1* – encoding a UVA/violet-light sensitive opsin (*19*, *20*) – acts in larval depth sensing (*21*), as well as in wavelength ratio detection involved in seasonal UV sensation in adult worms (*20*). *r-opsin*– and *c-opsin*-type *opsins* are generally thought to be expressed in distinct cell types, with the eyes of onychophorans, basally branching panarthropods, as possible exception of unknown meaning (*22*, *23*).

So far, evolutionary comparisons of *Platynereis* cell types have largely focused on larval stages, leaving open if there are aspects of adult neurogenesis and its regulation that are distinct or comparable between the bristleworm and representatives of other key phylogenetic groups. At the same time, the adult *Platynereis* brain appears as a suitable paradigm for investigating regulatory neurobiology. The brain is known to be the source of hormonal cues that orchestrate the major transition between regenerative, non-reproductive life stages (immature and premature stage) and a non-regenerative, reproductive stage (mature female and male worms) (*24–27*), which is accompanied by significant changes in the overall head transcriptome (*28*). Like in the bristleworm system, stem cell and regenerative potential in mammals decreases with reproductive maturity and age (*29*). By contrast, commonly studied invertebrate model systems like planarians or cnidarians do not exhibit a similar decline of their stem cell capacities (*30*, *31*).

Here, we characterize the molecular and cellular changes in the adult *Platynereis* head that accompany the transition between the animal’s major adult life stages. We find that the adult *Platynereis* brain harbors different classes of cells with a neurogenic potential, which exhibit marked similarities to active and quiescent mammalian aNSCs. Our investigation also leads us to the identification a *bona fide* stem cell system in the four camera-type eyes of the adult worms. Molecular markers and the position of these cells in the region adjacent to the glass body/lens draw a parallel to the ciliary marginal zone of vertebrate camera-type eyes. We also find the ciliary opsin gene *c-opsin1* to be co-expressed and functional in the early, proliferative stage of the rhabdomeric photoreceptor lineage.

## Results

### Adult brain neurogenesis is regulated during sexual maturation

In order to gain insight into molecularly defined cell types of the adult *Platynereis* brain, as well as their changes over maturation into the two sexes, we performed single-cell RNA sequencing (scRNA-seq) of worm heads of four distinct stages (immature, premature, mature female, mature male). After filtering and matching the sizes of the four libraries, our set comprised 19908 cells and 31811 reference gene sequences, with an average of 671 reference sequences expressed per cell.

Bulk RNA sequencing previously revealed significant changes in the total head transcriptome upon maturation (*28*). When integrating the individual libraries employing a merging approach, previously successfully used for developmental scRNA-seq time series (*32*), we obtained a representation in a uniform manifold approximation and projection (UMAP)-reduced space in which cell populations from immature and premature libraries grouped together, but separated from cell populations derived from mature female and male libraries (**Fig. 1A**). In contrast, data integration using the standard Seurat pipeline (*33*) resulted in a rather homogeneous distribution, lacking representation of the different adult stages (**fig. S1A**). Consistently, cluster identities obtained by basic clustering of the standard-integrated data were transferrable onto the merge-integrated data, where adjacent cell groupings in the merged data reflected subdivisions of the same transferred cluster identity (Materials and methods: Transfer of cluster identities between datasets of the same origin) (**Fig. 1B, C**).

**Figure 1:**
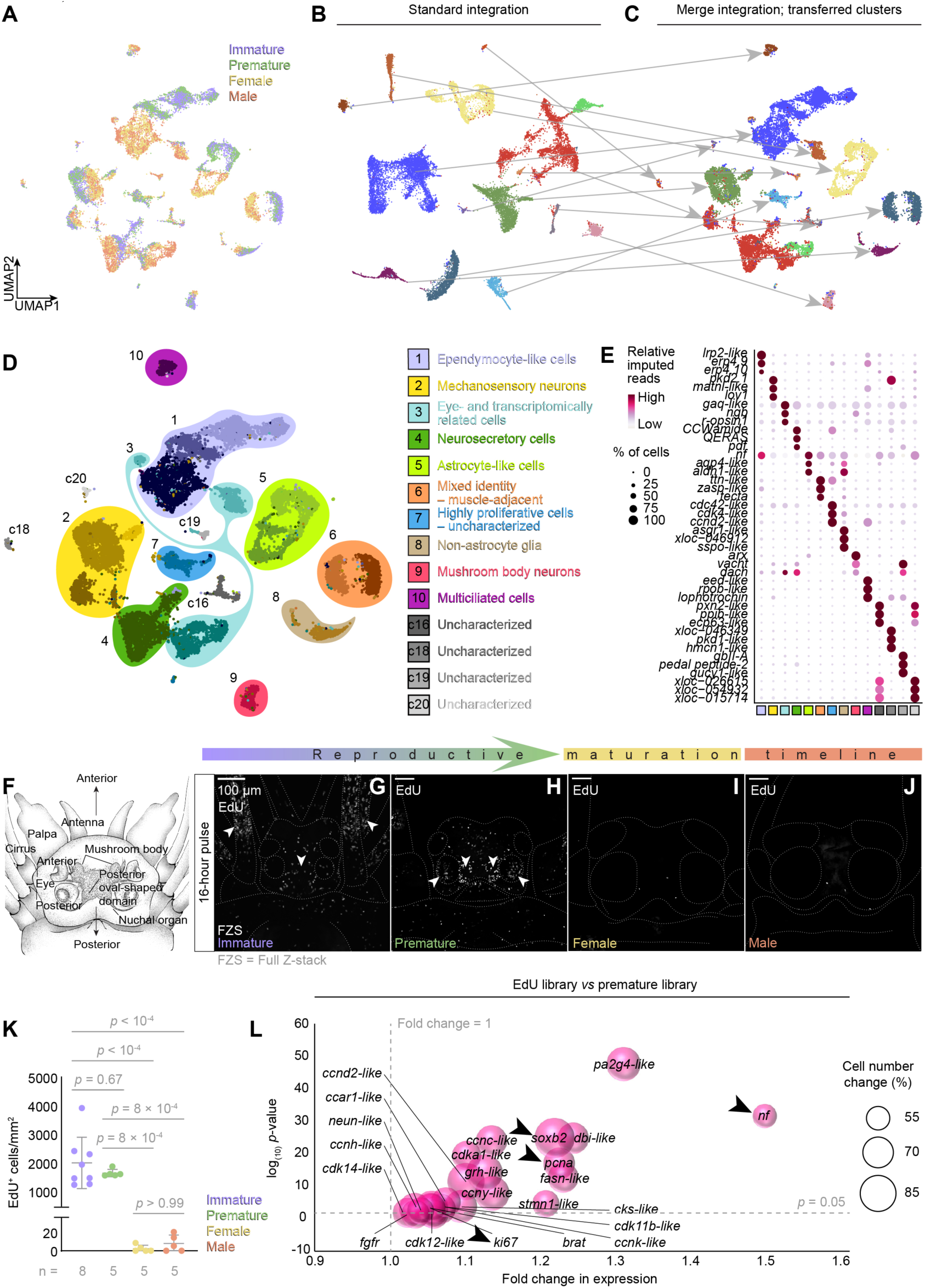
A single-cell transcriptomic atlas of the *Platynereis* brain and proliferative cell analyses reveal specific markers for cell types, including neural stem cells. **(A)** Merge-integrated immature, premature, female and male scRNA-seq libraries. **(B)** Basic clustering of standard-integrated libraries with **(C)** color-matched cluster identities transferred to the merge-integrated dataset. **(D)** Manually assigned annotation of overarching cluster identities (different color bubbles) consisting of one or more standard clusters (shades within bubbles) and **(E)** their respective marker genes. See fig. S1 for further details. **(F)** Schematic worm head. EdU incorporation detected in **(G)** immature, **(H)** premature, **(I)** female and **(J)** male worms following a 16-hour EdU pulse. Arrowheads indicate major mitotic foci; images are 72, 76, 36 and 36 µm thick, respectively. **(K)** Individual numbers of EdU-positive cells following a 16-hour pulse normalized to head area of immature (n = 8), premature, female and male worms (n = 5, respectively). One-way ANOVA with Tukey’s multiple comparisons test. **(L)** *Bona fide* neurogenesis- and proliferation-associated genes differentially expressed between whole EdU-labeled and -unlabeled premature libraries. Only genes surpassing the indicated fold change and p-value thresholds are displayed. Arrowheads indicate genes mentioned in main text; statistics by Wilcoxon Rank Sum test. Further details in **fig. S2**.

Our merge-integrated single-cell transcriptomic atlas thus reflects reproducible cell identity clusters and the broad transcriptomic changes between non-reproductive and reproductive stages of the aforementioned bulk transcriptomic analysis (*28*).

Annotation of the cell populations in the atlas, based on cluster markers and independently generated gene correlation modules (**fig. S1B-N; Auxiliary file Tab 2**), resulted in 10 broad categories of cell identities and 4 smaller populations (**Fig. 1D, E**; **Auxiliary file Tab 1)**. These included likely neurosecretory, photo- and mechanosensory populations, as well as cells corresponding to brain structures like mushroom bodies. We also identified several *bona fide* glial cell populations, including astrocyte-like cells, non-astrocyte glia and ependymocyte-like cells, as well as cells of combined neuronal and non-neuronal signatures, such as multiciliated cells, muscle-adjacent cells and a highly proliferative uncharacterized cell population (**Fig. 1D, E, Auxiliary file Tab 1**).

Given the presence of cells characterized as proliferative by molecular signatures and in order to better understand the possible plasticity of these cell type clusters across the same adult stages, we probed for cell proliferation in the adult brain by using 5-ethynyl-2’-deoxyuridine (EdU) incorporation, which demarcates S-phase nuclei. Heads of non-reproductive adult animals exhibited high levels of proliferation (**Fig. 1F-H, K**, **Auxiliary file Tab 3**). By contrast, heads of reproductive worms showed nearly no EdU-positive (EdU^+^) cells (**Fig. 1F, I-K**, **Auxiliary file Tab 3**).

To assess whether the EdU^+^ cells belonged to neurogenic lineages, we introduced an EdU labeling step into the scRNA-seq protocol and used fluorescence-activated cell sorting (FACS) to isolate and sequence EdU^+^ cells from premature heads. The obtained scRNA-seq transcriptomes of EdU^+^ cells clustered with immature and premature cells in the UMAP representation of the library projected over the UMAP-reduced space of the brain atlas (Materials and methods: Predictive dimensional reduction and annotation of new datasets), matching their origin (**fig. S2A, B**: henceforth called ‘predicted UMAP projection’). Compared to unlabeled cells, EdU^+^ cells not only showed significantly enriched expression of proliferative markers and cell cycle genes such as *proliferative cell nuclear antigen*/*pcna*, *cyclin-dependent kinase 1-like*/*cdk1-like* (**Fig. 1L**, **fig. S2C**), but also exhibited an enrichment for an established marker of *Platynereis* neurogenesis, *soxB2* (*34*, *35*) (originally referred to as *soxB*), as well as the previously uncharacterized orthologs of mammalian *ki67* and *gfap* (here referred to as *neurofilament*/*nf*) that are known to be important markers of proliferative and undifferentiated cells with neurogenic potential in mammals, respectively (arrows in **Fig. 1L**, **fig. S2C**, **fig. S3A-C, Auxiliary file Tab 4**). Ki-67 orthologs have not yet been reported in other invertebrates. Our annotation is supported by phylogeny and protein domain analyses (**fig. S3B, C**). The latter revealed both the N-terminal Forkhead associated domain (FHA) and the protein phosphatase 1 (PP1) binding domain diagnostic of vertebrate Ki-67 members (**fig. S3C**). PP1 is known to be relevant for the function of Ki-67 in mitotic progression (*36*). Moreover, the predicted bristleworm Ki-67 protein has a content of serine/threonine residues of 20%; akin to mouse Ki-67, where these residues are targets of cell-cycle-dependent phosphorylation (*36*). Taken together, scRNA-seq of the EdU^+^ cells identified several neurogenic markers, pointing at molecular similarities between the aNSCs present in the bristleworm and in the mammalian brain.

### A ciliary marginal zone-like stem cell system in the adult eye generates eye photoreceptors and support cells

The discovery of aNSC-like cells in the *Platynereis* brain led us test if there were specific neurogenic domains contributing to adult eye and brain growth. Specifically, we investigated the adult eyes of the animals that exhibit an everted, camera-type setup (*37*). The eyes show continuous growth, ending with a brief period of rapid augmentation shortly prior to the animals reaching sexual maturity (**Fig. 2A-D**). It has been hypothesized that the eyes generally grow by continuous apposition of cells to the rim of the retina, while the final eye enlargement was thought to be caused by expansion of the glass body/lens volume, without further cell number increase (*37*). While we confirmed a marked increase in glass body/lens diameter in reproductive worms (**fig. S4A-E**), our EdU single-cell analyses suggested the continuous presence of neurogenic cell populations in this organ (**fig. S2A, B)**. On the cell type level, we therefore focused on population 3 of our transcriptomic map (**Fig. 1D, E**; **Fig. 2E**; eye- and transcriptomically related cells) that is characterized by a cumulative module built from genes previously identified as expressed in *r-opsin1*-positive photoreceptor cells (*38*), as well as on an overlapping, independently generated gene correlation module (**fig. S4F-H**; **fig. S1H**, **Auxiliary file Tab 5, Auxiliary file Tab 2 (Module 7)**). In contrast to the notion that proliferation played no significant role in eye growth during maturation, the contribution of cell libraries derived from reproductive animals was approximately fourfold higher than those derived from non-reproductive stages, with males exhibiting an even higher cell count than females (population 3 in **fig. S4I**), consistent with their final eye-to-head size (**Fig. 2C, D**, **fig****. S4J**, **Auxiliary file Tab 3**).

**Figure 2.**
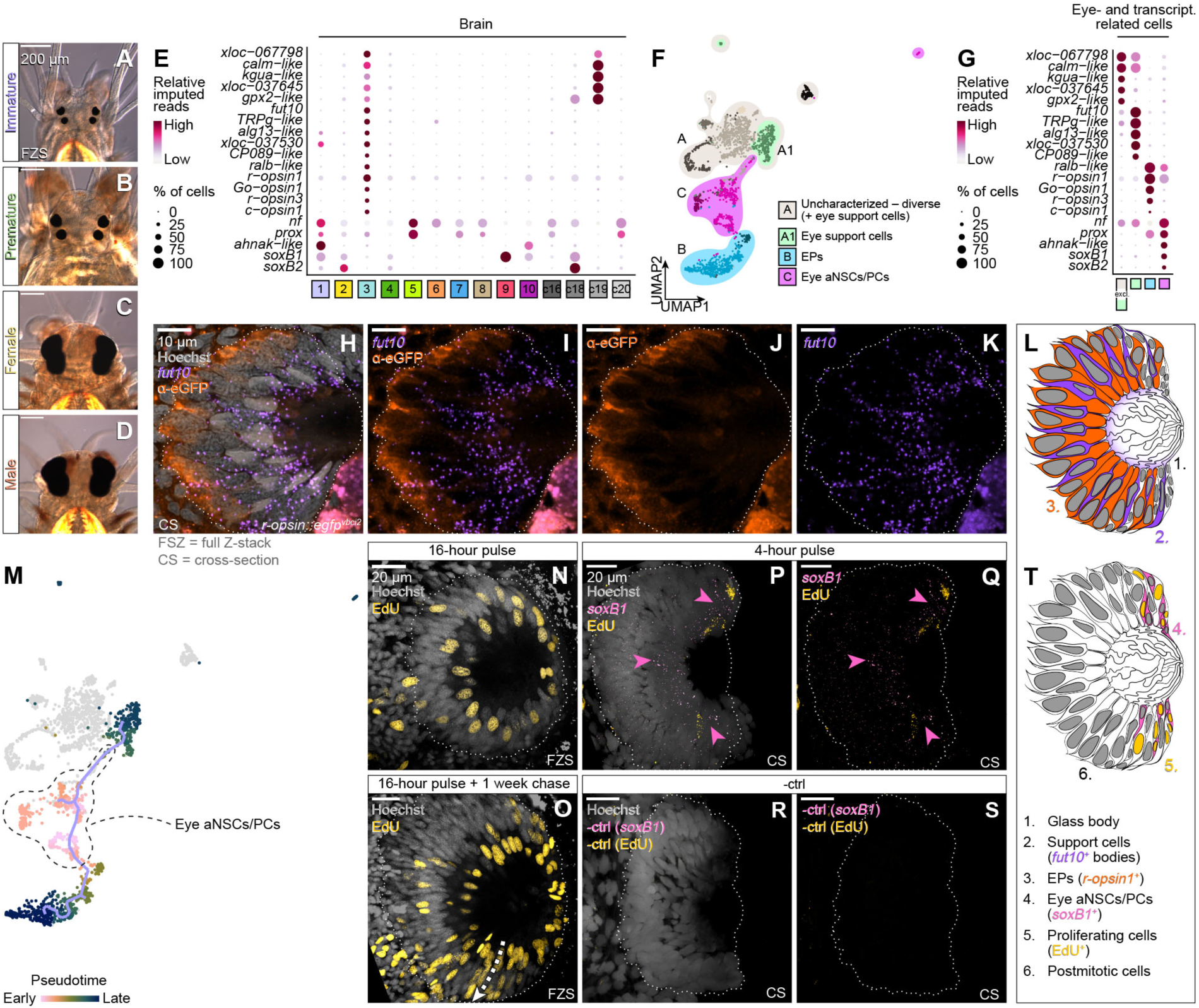
The adult eye grows via a CMZ-like zone, giving rise to support– and photoreceptor cell lineages. Eye size comparison in differential interference contrast micrographs of **(A)** immature, **(B)** premature, **(C)** female and **(D)** male worm of 15, 15, 49 and 63 µm thickness, respectively. **(E)** Markers of eye- and transcriptomically related cells in the brain atlas. **(F)** Subclustered eye- and transcriptomically related cell dataset, colored by manually assigned overarching cluster identities (bubbles) and standard clusters (cells colored in different shades within a bubble); eye support cells cluster together with the rest of the population in the upper partition of the UMAP. **(G)** Genes from **(E)** in dataset from **(F)**. **(H-K)** HCR staining of *fut10* and immunofluorescent labeling of eGFP in an *r-opsin1::egfp^vbci2^* worm. Thickness: 1.15 µm; signal in lower right corner below eye is autofluorescence. **(L)** Schematic representation of **(H)** as a medial cross-section of the eye. **(M)** Putative differentiation trajectory of cells corresponding to the eye lineages ordered in pseudotime according to inferred differentiation states; cells in grey are suspected brain cells with a transcriptomically similar signature. EdU detection following a **(N)** 16-hour pulse and **(O)** additional 1-week chase period. Arrow indicates suggested direction of lineage progression; thickness: 29.01 and 13.04 µm, respectively. **(P-S)** HCR labeling of *soxB1* and EdU incorporation following a 4-hour pulse and the corresponding negative control (-ctrl); arrowheads indicate expression regions of *soxB1*; both images 9.41 µm thick. **(T)** Schematic representation of **(P)** as a medial cross-section of the eye. Further details in **fig. S4-6.**

To obtain a better understanding of the possible neurogenic processes contributing to eye growth, we subclustered the cells contributing to population 3 into a dataset of 2324 cells (immature: 238, premature: 275, female: 672, male: 1139). When we plotted the expression of the main photoreceptor molecule of the adult eyes, *r-opsin1* (*18*, *38*) onto the subclustered dataset (**fig. S5A**), the cells split into three main subclusters: an *r-opsin1*-negative partition (A), a strong *r-opsin1*-positive partition (B), and a smaller partition positioned between these with weaker and patchier *r-opsin1* expression (C) (**Fig. 2F, fig. S5A**). Aiding this annotation, we also identified distinct genes enriched in each of the populations and plotted them in both the eye and the brain dataset (**Fig. 2E, G**; **Auxiliary file Tab 6**).

Several genes associated with photoreception and -transduction demarcate the *r-opsin1*-positive partition B, confirming them to be eye photoreceptors (EPs) (**Fig. 2E, G**; and see below). Based on past ultrastructural work, a major cell type of the *Platynereis* eye besides the EPs are support cells, with roles in formation of the glass body/lens and pigmentation (*37*, *39*). A subset of partition A showed particular enrichment for *fut10* (**fig. S5B**), the *Platynereis* ortholog of α-1,3-Fucosyltransferase 10 (*40*) (**fig. S5C**). *In situ* HCR analysis showed the strongest expression of *fut10* enriched in cells that match eye support cells by frequency and position (**Fig. 2H-L, fig. S5D-F**). Outside the eye, *fut10* also shows some expression in the posterior oval-shaped domains of the brain (**fig. S5D-F**). Based on our stainings, we designated a subset of partition A as eye support cells (A1 in **Fig. 2F, G**), while the remaining section of partition A (here denominated as “uncharacterized – diverse”) likely contributed to photoreceptors and pigment cells known to be present outside the eye, in the worms’ brains (*17*, *41*, *42*). Whereas these are not the only brain photoreceptors (**fig. S7A-Q**), they are molecularly closely related to the eye photoreceptors.

Partition C is the group of cells interconnecting support cells (partition A1) and photoreceptor cells (partition B) (**Fig. 2F**). Cells in this partition expressed neurogenic markers like *soxB1, soxB2, prox* and *nf* (**Fig. 1L**, **Fig. 2F, G, fig. S2C, fig. S6A-C; Table S1**). Consistently, differentiation trajectories inferred by Monocle3 (*43*) suggested these cells as possible origin of *r-opsin1^+^* eye photoreceptors (partition B) and *fut10*^+^ eye support cells (partition A1) (**Fig. 2M, fig. S6D**). Furthermore, both the cumulative module computed from transcripts enriched in the EdU^+^ cells in the eye- and transcriptomically related cell dataset, as well as a predicted UMAP projection of EdU^+^ cell transcriptomes in the dataset populated the trajectories of this partition towards partitions B and A1 (**fig. S6E, F**). This is consistent with the four day-long EdU pulse the animals were exposed to prior to sequencing, hence the majority of the EdU^+^ cells likely belong to a population of actively dividing cells on their path towards differentiation. We thus reasoned that this partition contained stem cells supplying the eye photoreceptor and eye support cell populations, and henceforth refer to partition C as eye adult neural stem/progenitor cells (eye aNSCs/PCs) (**Fig. 2F**).

We next wondered whether the eye aNSCs/PCs reside in a confined area of the eye, or are spread throughout the eye cup. For this, we performed additional, shorter EdU uptake assays with and without a chase period. In the pulse-only experiment, the majority of EdU^+^ cells localized to the innermost cell ring of cells surrounding the glass body/lens, appearing to nest between photoreceptor rhabdomes and support cell processes, with a few cells located more distally to the glass body/lens (**Fig. 2N**). Following a long chase period, the number of EdU^+^ cells in the innermost ring multiplied, and we observed trails of EdU^+^ cells arching outwards (arrow in **Fig. 2O**), reminiscent of the spreading of fish retinal clones along arched continuous stripes (*44*, *45*). Finally, we employed a combined approach of *in situ* HCR and EdU labeling to probe for a link between the proliferative zone in the eye and the *in silico*-characterized eye aNSCs/PCs. Indeed, *soxB1*, the prominent marker of both vertebrate retinal stem cells and eye aNSCs/PCs, delineated the innermost cell layer in the eye, where most of the early EdU incorporation took place (arrowheads **Fig. 2P-T**). In summary, these data support the presence of a stem cell zone at the edge of the retina abutting the glass body/lens, giving rise to new eye photoreceptors and eye support cells.

### The ciliary opsin *c-opsin1* is required for proper differentiation of *r-opsin1*-positive photoreceptors

The transcriptomic characterization of eye photoreceptor cells not only confirmed the presence of the r-opsin genes *r-opsin1* and *r-opsin3* as well as the G_o_-opsin gene *G_o_-opsin1* in the adult eyes, but also indicated expression of the c-opsin gene *c-opsin1* (**Fig. 2G**, **fig. S7A, B, E, G**). While *r-opsin1, r-opsin3* and *G_o_-opsin1* had been previously described in the eyes (*38*, *41*, *46*–*48*), *c-opsin1* was so far only known for its expression in the larval brain, where it exclusively demarcates extraocular non-visual photoreceptor cells (*17*). Visualization of both *r-opsin1* and *c-opsin1* in the subclustered eye- and transcriptomically related cell transcriptome map indicated that *c-opsin1* was co-expressed with *r-opsin1* in a distinct subpopulation of cells that reside on the transition between the aNSCs/PCs and the start of the photoreceptor differentiation trajectory (**Fig. 3A-D**).

**Figure 3.**
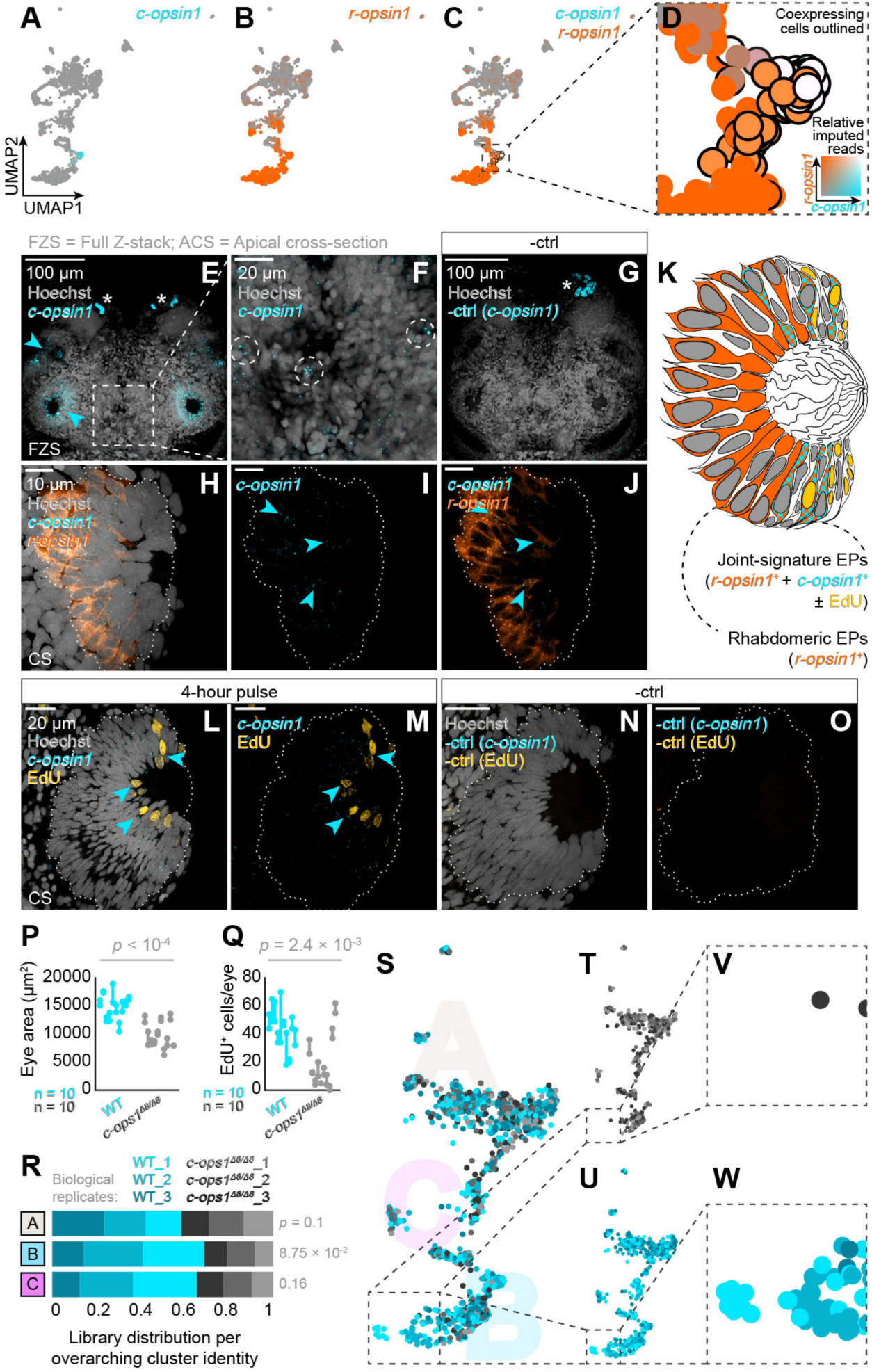
A ciliary opsin-gated system helps modulate rhabdomeric photoreceptor differentiation. **(A-D)** Co-expression of *c-opsin1*and *r-opsin1* in the eye predicted by scRNA-seq. **(E-G)** HCR detection of *c-opsin1* in the head and the corresponding negative control; circles indicate brain staining. Images are 56.66, 56.66 and 48.46 µm thick, respectively. Asterisks: autofluorescent glands. **(H-J)** HCR co-detection (arrowheads) of *c-opsin1* and *r-opsin1* in the eye; thickness: 1.15 µm. **(K)** Schematic representation of **(H)** as medial cross-section of the eye. **(L-O)** HCR labeling of *c-opsin1* and EdU incorporation in the eye following a 4-hour pulse and the corresponding negative control. Thickness: 1.15 and 2.58 µm, respectively; arrowheads: double-positive cells. Nested scatterplots of two posterior eyes per animal comparing **(P)** eye size and **(Q)** EdU-positive cell count following a 4-hour pulse in WT (n = 10) and *c-ops1^Δ8/Δ8^*(n = 10) worms; statistics: nested t-test. **(R)** Representation of WT and *c-ops1^Δ8/Δ8^* scRNA-seq libraries (n = 3 for each genotype) in overarching cluster identities of the eye- and transcriptomically related cell dataset; statistics: unpaired t-test. **(S-W)** WT and *c-ops1^Δ8/Δ8^* libraries in UMAP-reduced space of existing eye- and transcriptomically related cell dataset. Further details in **fig. S8.**

To probe for the validity of these data, we performed *in situ* HCR analyses on non-reproductive worms’ head and eye. Detection of *c-opsin1* revealed a few individual cells in the posterior oval-shaped domains of the brain (dotted circles **Fig. 3E-G**), consistent with the aforementioned larval expression data (*17*). However, the most prominent *c-opsin1* expression indeed occurred in cells within both pairs of adult eyes (arrowheads **Fig. 3E-G**). Co-detection of *c-opsin1* and *r-opsin1* mRNA confirmed that these cells co-express both of the opsin genes (**Fig. 3H-K**). The position of such cells adjacent to the glass body/lens, where the aforementioned results revealed the CMZ-like stem cell zone (**Fig. 2T**, **Fig. 3K**), suggests that that they are part of the predicted aNSCs/PCs on their differentiation trajectory towards eye photoreceptors (**Fig. 2M**, **Fig. 3A-D**). To verify this hypothesis, we also investigated if any of the *c-opsin1-*positive cells were proliferative. Indeed, detection of *c-opsin1* RNA in individuals labeled by a short EdU pulse revealed that a subset of the EdU^+^ cells close to the glass body/lens indeed expressed *c-opsin1* (**Fig. 3L-O)**, suggesting that *c-opsin1*^+^ cells were not terminally differentiated, but demarcate a transitory state that gives rise to the *r-opsin1^+^* photoreceptor population of the adult eye.

If *c-opsin1* plays a role in the differentiation of *r-opsin1*-positive photoreceptors, we reasoned that genetic ablation of the *c-opsin1* gene could have an impact on the development and/or molecular signature of eye photoreceptors. To test this idea, we first compared the size and proliferation in eyes in wild-type (WT) animals with individuals with homozygous mutations in the *c-opsin1* gene (*c-ops1^Δ8/Δ8^*(*20*)). As quantified by their area, eyes of *c-ops1^Δ8/Δ8^* individuals were consistently smaller than their WT counterparts (**Fig. 3P; Auxiliary file Tab 3**). Likewise, EdU labeling revealed a smaller number of proliferative cells in eyes of *c-ops1^Δ8/Δ8^* individuals (**Fig. 3Q; Auxiliary file Tab 3**).

In order to assess possible changes in the transcriptomically defined cell populations of the eyes, we finally performed a scRNA-seq experiment on premature *c-ops1^Δ8/Δ8^* individuals and their WT counterparts. We predicted UMAP projections of the resulting *c-ops1^Δ8/Δ8^* and WT transcriptomes in the UMAP-reduced space of the brain atlas and the eye- and transcriptomically related cell subset (**fig. S8A-D**). Contribution of cells from libraries generated from *c-ops1^Δ8/Δ8^* specimens *versus* those generated from WT controls to the different clusters differed significantly in five cases, in which four showed a significant underrepresentation (**fig. S8E**).

Among those is the eye- and transcriptomically related cell overarching cluster identity, with a notable reduction in the number of cells in the EP population (**Fig. 3R, fig. S8E)**. While such an analysis represents the impact of the mutation over the entire life of the animal and hence may show effects that are also outside of the *c-opsin1* expressing cells themselves, this finding is in line with the observed reduction of eye size in mutants and the hypothesis that *c-opsin1* influences the division rate and/or differentiation of *r-opsin1^+^* eye cells.

Indeed, when assessing the specific UMAP cell coordinates of transcriptomes from both genotypes, eye photoreceptor cells from the *c-ops1^Δ8/Δ8^* individuals lacked a group of cells at the end of the cluster in both the standard and predicted UMAP-reduced spaces (**Fig. 3R-W**; **fig. S8F, G, fig. S13A-C**). Taken together, these data are consistent with a function of *c-opsin1* in modulating the differentiation of *r-opsin1*^+^ eye photoreceptors.

### Reproductive maturation imposes a special cell state combining neurogenic signatures with markers of mammalian aNSC quiescence

In order to better further understand the eye and brain aNSC/PC plasticity, we finally asked which changes these cells undergo when the animals enter their reproductive stage, at which EdU incorporation is basically absent (**Fig. 1F**, **Fig. 1I-K**).

In order to test for molecular signatures distinguishing cells before and after maturation, we performed a pseudo-bulk differential expression analysis between all cells from non-reproductive and reproductive stages (**Auxiliary file Tab 7**). We then conducted gene ontology (GO) analyses on the sets of differentially regulated genes to determine the most distinct biological processes for each set. In non-reproductive, proliferative worms, the top enriched GO terms were related to translation, ribosomal processes, as well as coding and non-coding RNA processing (**Fig. 4A**, **Auxiliary file Tab 8**). In reproductive, non-proliferative worms, top enriched terms included intracellular signaling and synaptic and vesicular processes (**Fig. 4B**, **Auxiliary file Tab 9**). These, as well as other enriched terms from non-reproductive (blue stars in **Fig. 4C**) and reproductive (orange stars in **Fig. 4C**) gene sets, resembled signatures associated with active and quiescent mammalian aNSCs, respectively (*11*, *49*, *50*) **(Fig. 4C)**. In contrast, no GO terms associated with senescence or cell death were enriched in the cells from reproductive (or non-reproductive) animal heads, indicating that the non-proliferative state in the reproductive worms was likely not the result of an irreversible cell cycle arrest, apoptosis or necrosis (**Auxiliary file Tab 8, 9)**. While the association of mammalian quiescence hallmarks with genes enriched in cells from reproductive animals was generally true, we noted a few exceptions for terms related to proliferation and neurogenesis, which were significantly overrepresented in supposedly non-proliferative reproductive animals (**Auxiliary file Tab 9).** Specifically, we observed that transcripts contributing to these terms were genes associated with neuronal proliferation in progenitor cells, like *prospero/prox*, *soxC* (ortholog of vertebrate *sox4*, *sox11* and *sox12*) and *par3a* (ortholog of vertebrate *pard3*) (*34*, *51*). While other genes associated with active aNSCs/PCs remained expressed in the reproductive stage, *prox*, *soxC* and *par3a* were even upregulated in reproductive animals (**Fig. 4D-I, fig. S11A-F**).

**Figure 4.**
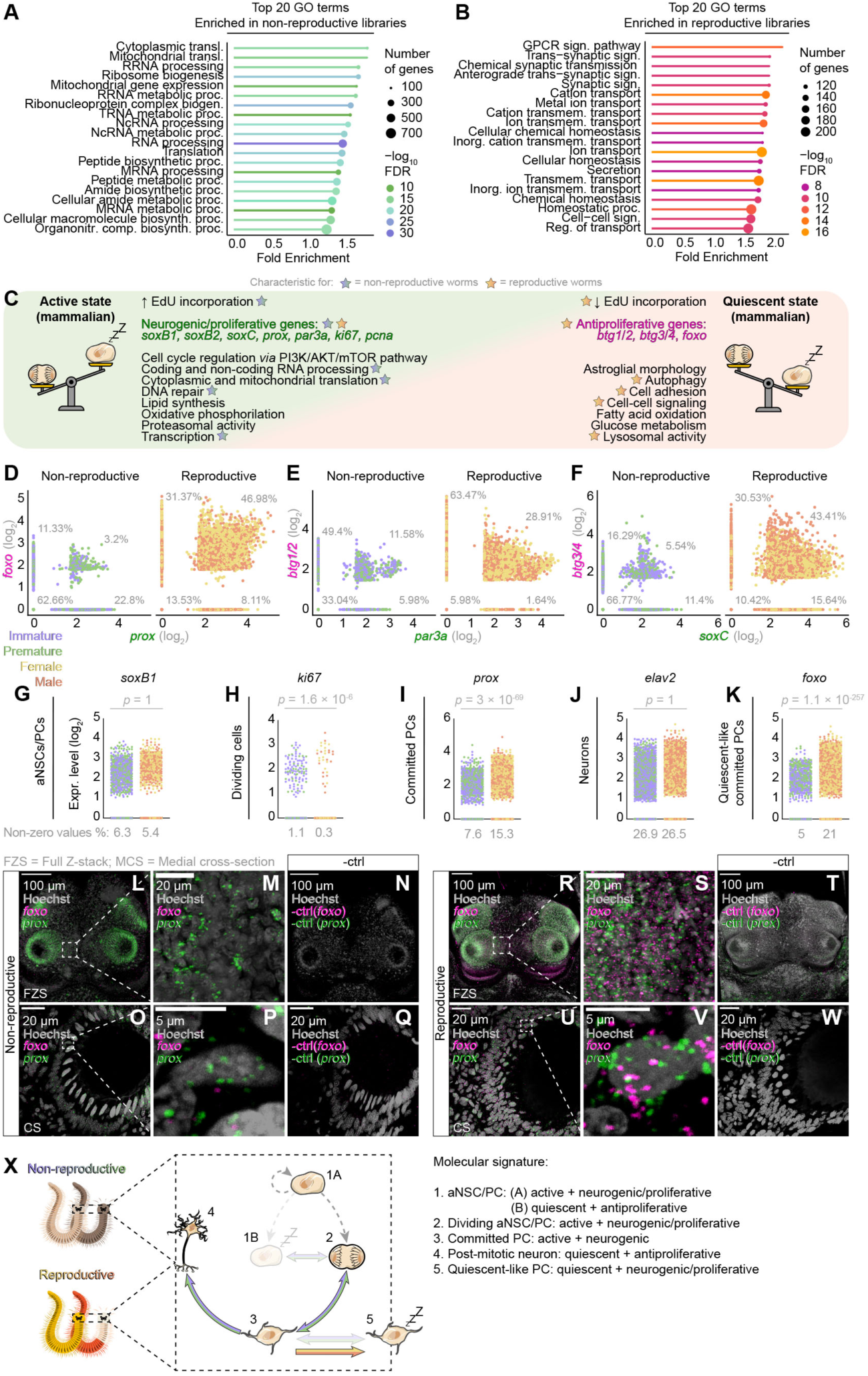
Vertebrate-type neurogenic and quiescent-like signatures are co-regulated in reproductive brains. Top 20 GO terms associated with genes enriched in **(A)** non-reproductive and **(B)** reproductive libraries; FDR = false discovery rate. **(C)** Hallmarks of active and quiescent aNSCs in mammalian systems; stars indicate physiological and transcriptomic hallmarks detected in non-reproductive (blue stars) and reproductive libraries (orange stars) in the GO term analysis of the genes differentially expressed between the libraries. **(D-F)** Relative imputed values of co-expression of neurogenic (green) and quiescence-associated genes (magenta) in non-reproductive and reproductive libraries; percentages indicate fraction of cells expressing one, both or none of the markers. **(G-K)** Scatterplots showing non-imputed expression of representative genes characteristic for distinct stages of the neurogenic cascade in non-reproductive and reproductive libraries. Statistical significance calculated using a Wilcoxon Rank Sum test. HCR co-labeling of *foxo* and *prox* and corresponding negative controls in a **(L-N)** premature head (55.17, 55.17 and 66.35 µm thickness, respectively), **(O-Q)** premature eye (2, 2 and 1.98 µm respective thickness), **(R-T)** mature male head (94.69 µm thickness in all images) and **(U-W)** mature male eye (2 µm thickness in all images). **(X)** Schematic representation of the proposed progression of the neurogenic cascade in non-reproductive and reproductive animals. Further details in **fig. S11**.

However, consistent with the absence of cell divisions, we also found enrichment of genes associated with the quiescent stem cell state in the same gene set, such as orthologs of the mammalian genes *B cell translocation gene 1/btg1*, *btg3*, *foxo3* and *foxo6,* which we identified as *Platynereis btg1/2*, *btg3/4* and *foxo*, respectively. Finally, we detected unchanged expression levels of post-mitotic markers *synaptotagmin/syt* and *embryonic lethal/abnormal vision* genes *elav1* and *elav2*, indicating that the switch in neurogenic capacity does not influence the ratio of resulting neurons (**Fig. 4D-F, J, K, fig. S9, fig. S10, fig. S11G-J**). FoxO proteins are relevant for regulating stem cell quiescence in diverse stem cells (*52*, *53*). Btg1 and 2 have been shown to stabilize mammalian stem cell quiescence by reducing polyadenylation of mRNAs (*54*), while Elav and Syt are involved in neuronal differentiation (*55*) and neuronal function (*56*).

The simultaneous co-enrichment of neurogenic and antiproliferative genes in cell signatures of reproductive animals suggests that the cessation of cell divisions is linked to the induction of a quiescence-like molecular program in the formerly proliferative cells of the brain and eye of the reproductively maturing animals. Bioinformatic approaches using cumulative gene modules computed for *bona fide* markers of distinct phases of neurogenesis as well as quiescence-associated genes confirmed a high overlap in cells expressing neurogenic and quiescence-associated markers in reproductive animals (**fig. S11B, D, F, H, J**).

To experimentally validate this finding, we performed HCR of representative markers on non-reproductive and reproductive animals (*57*). Animals of both stages displayed expression of *prox*, concentrating in the eyes and posterior oval-shaped domains of the brain (**Fig. 4L-Q**). By contrast, and in agreement with the *in silico* data, the expression of *foxo* was markedly enriched in reproductive worms, coinciding with *prox* in the same cells (**Fig. 4R-W**).

Taken together our data suggest that reproductive maturation imposes a quiescence-type signature on formerly proliferative cells, resulting in a sharp downregulation of proliferation at this stage, despite the simultaneous presence of a molecular signature associated with neurogenesis (**Fig. 4C, X).**

## Discussion

Comparative model systems are valuable sources for neurobiological discovery and mechanistic principles (*58*). In line with this idea, *Platynereis dumerilii* has served as a reference model for comparing both early nervous system patterning and cell types across bilaterians (*17*, *59–62*). So far, most emphasis has been on larval development, building on the idea that the trochophore larva maximizes comparability across different phyla (*63*), and focusing on functional differentiated cell types as units of comparisons (*60*, *64*). By extending the analyses to adult stages, our work allows us to explore signatures of adult neural stem cells and their regulation in the head – processes that are neither accessible in larvae, nor well understood in their evolutionary history. As outlined, our work reveals modulation of neurogenesis on two levels: first, the growth of the adult camera-type eye *via* a circular stem cell zone, reminiscent of the vertebrate ciliary marginal zone, and coupled to the unexpected expression of the ciliary opsin gene *c-opsin1*, modulating rhabdomeric photoreceptor cell differentiation. Second, a global molecular signature reminiscent of mammalian aNSC quiescence that is imposed by the entrance of animals into the reproductive stage and is associated with a global stop of brain proliferation.

The bristleworm aNSC system exhibits both similarities and differences to mammalian aNSCs. Molecularly, the *gfap* and *ki67* orthologs we find in the worm, and their association with *bona fide* aNSCs, support the notion that there is an ancient molecular signature of neurogenic/ proliferative brain cells hat is shared between bristleworms and vertebrates (*34*). The bristleworm Ki-67 ortholog we report here dates the emergence of this regulator to early bilaterian ancestors. In contrast to mammalian aNSCs, however, where quiescence is best studied as a mechanism regulating the activation/inactivation of an early neuronal stem cell pool, we find the quiescence-like signature in bristleworms to be mostly associated with progenitor cells that stop proliferating in reproductive maturity. It suggests that this association relates to a more general, anti-proliferative function of the signature in worms. Consistently, at least some vertebrate orthologs like Btg2 also have an anti-proliferative role in progenitor cells undergoing terminal neuronal differentiation (*65*).

Our study has not attempted to address if or not the quiescence-like state of worm aNSCs/ PCs is reversible. Like other nereidid annelids, *Platynereis* exhibits semelparity, with mature animals dying after reproduction, such that post-reproductive individuals cannot be tested for reactivation of neurogenesis. However, this semelparous reproductive mode likely represents a secondary evolutionary modification, as related worms have been demonstrated to possess a latent capacity for iteroparity, *i.e.* multiple reproductive cycles within lifetime (*68*). Moreover, representatives of other annelid groups such as the Palolo worm *Eunice viridis* only transform posterior segments during reproduction, and thus exhibit a natural cycle of reproductive and regenerative states (*69*). In this setting, a temporary quiescence of neurogenesis may serve as a way of directing energy expenditure towards gametogenesis, consistent with the idea that in organisms, quiescence is a mechanism to control energy consumption by growth (*70*).

Cephalopod camera-type eyes have served as text book examples of convergent morphological evolution. Like cephalopods, *Platynereis* worms exhibit everted camera-type eyes. Our molecular characterization of cell populations in the growing bristleworm eye exhibits some parallels to gene expression data recently described for the developing squid retina (*71*). For instance, *soxb1* demarcates early progenitor cells in both systems. Also, *soxB2* gene is expressed in the progenitor zone of both systems, even though our data suggest this to be more related to the support cell lineage than the eye photoreceptor lineage. Whereas a spatially defined growth zone has not been reported for cephalopod eyes, the spatial gene expression data we report here suggest the existence of a dedicated stem cell zone in the bristleworm eye. By the position of these cells close to the glass body/lens and by their likely contribution to both pigmented support cells and photoreceptors, we suggest this zone to be a functional correlate of the vertebrate ciliary marginal zone, extending a functional building principle in camera-type eye analogous evolution to the stem cell zone.

The finding that the ciliary opsin *c-opsin1* impacts on the proper differentiation of the *r-opsin1*-positive photoreceptor lineage adds to our understanding of photoreceptor evolution. Whereas the evolutionary divergence between distinct *r-opsin* and *c-opsin* genes dates back to pre-bilaterian times (*17*), these receptors are closest neighbors in deep phylogenetic analyses (*72*), suggesting that they duplicated from a joint precursor. This scenario implies that, before the emergence of distinct photoreceptor subtypes expressing any single of these *opsin* genes, they would both have been co-expressed in the same cell type, allowing for functional diversification.

The situation we observe in today’s bristleworm eyes might thus either indicate a more recent co-option of *c-opsin1* for the regulation of the *r-opsin*^+^ eye photoreceptor lineage, or possibly reflect an ancestral constellation. The role of *c-opsin1* to modulate specific cell states and/or cell type of *r-opsin*^+^ eye photoreceptors might help to optimize the worms’ sensory system to the different light conditions they encounter, depending on settling habitat and season. Light in the marine environment provides organisms with a wealth of information (*73*). Adjusting growth processes of sensory systems might hence provide organisms with a level of plasticity that is a selective advantage in constantly changing surroundings. If such plasticity occurs in other animal groups, and if so, how it copes with anthropogenic changes, such as artificial light, will be important questions for future work.

### Materials and methods

#### Worm culture and light conditions

*Platynereis dumerilii* worm culture was maintained at the Marine Facility of the University of Vienna, Austria under a 16:8 light:dark regime with a cyclical nocturnal illumination simulating a full moon for 8 nights per one lunar cycle (*74*). For experiments involving *c-ops1^Δ8/Δ8^* animals and corresponding WT individuals, the animals were kept under full-spectrum sunlight conditions, as *c-opsin1* encodes a UV-sensitive photoreceptor (*20*, *75*); otherwise, the animals were reared in artificial sunlight conditions. All worms received artificial moonlight. Artificial sunlight and moonlight spectra:(*74*). Animals were raised in rectangular plastic containers filled with 1-1.5 L of 1:1 mix of sterile-filtered, salinity-adjusted natural and artificial sea water (Tropic Marin Classic). Batches were fed once a week with minced organic spinach leaves and once a week with a blend of spirulina powder and ground Tetramin fish food.

#### Worm staging and sampling

Experiments on animals were conducted following the Austrian and European guidelines for animal research. WT (PIN strain), *c-ops1^Δ8/Δ8^*(*20*) and *r-opsin::egfp^vbci2^* transgenic (*47*) worms were sampled between Zeitgeber Time (ZT) 2 and 3, a week following the full moon stimulus, unless otherwise indicated. Immature worms were chosen based on age (under 3 months) and size (30-40 segments). Premature worms were distinguished from immature worms by size (over 60 segments) and presence of developing gametes, microscopically detectable upon clipping the worm’s body between two posterior parapodia. Premature animals with an evident onset of metamorphosis (eye enlargement, emptied gut, color change from brown/grey to light orange, morphological changes in the first parapodia) were omitted from the study. Mature animals were selected for enlarged eyes, sexually dimorphic coloration (females: bright yellow; males: whitish anterior and bright red posterior) and crawling behavior. Fully mature animals exhibiting swimming behavior were omitted from the study due to the risk of spontaneous reproduction during handling. For scRNA-seq experiments involving EdU-treated worms, prior to sampling, premature animals were exposed to a 96-hour pulse of EdU, as described in “EdU labeling”; for these experiments, the treated and the control group (“sampling time-matched premature library”) were sampled at the time of new moon, i.e. two weeks following the full moon stimulus. All worm heads were sampled by anesthetizing the animals in 7.5% (w/v) MgCl2 diluted 1:1 in worm culture water, followed by a surgical amputation of the head. For scRNA-seq experiments, 20 immature, 10 premature, 8 female and 8 male worm heads (transcriptomic atlas), 120 EdU-treated premature heads (EdU experiment) and 10 premature WT and *c-ops1^Δ8/Δ8^* worms were collected per replicate (*c-opsin1* experiment) or sample (all other experiments). The heads were amputated anteriorly to the first segment and posteriorly to the nuchal organ commissure. For imaging experiments, heads were amputated posterior to the jaws. Unless otherwise indicated, all imaging experiments were conducted on premature worms as representative of the non-reproductive stage, and mature male worms as representative of the reproductive stage.

#### Generation of single-cell suspensions

Following the amputation of the worm heads, the tissue was simultaneously dissociated, permeabilized and fixed following the acetic acid-methanol (ACME) protocol (*76*) with the following modifications. Heads were incubated for 1 h in the ACME solution prepared according to the protocol, and tissue dissociation was mechanically aided by resuspending the solution for 2 min every 15 min. The suspension was filtered through a 40 µm cell strainer (Flowmi™ Cell Strainers; Scienceware) and centrifuged at 2000 × g, and the supernatant was discarded. Where applicable, the EdU Click-iT™ reaction was performed on EdU-treated samples for 20 min, according to the manufacturer’s protocol for the Click-iT™ EdU Cell Proliferation Kit for Imaging (Thermo Fisher Scientific); the sample was then centrifuged and the supernatant discarded. The pellets of all samples were suspended in the freezing solution prepared according to the protocol, using 0.1% instead of 1% BSA, and stored at -20°C. On the day of library preparation, the samples were thawed, stained for 7 min with 1 µg/mL nuclear dye (Hoechst 33342, trihydrochloride, trihydrate; Thermo Fisher Scientific) and centrifuged, the supernatant discarded, the pellet suspended in resuspension solution and transferred into tubes for fluorescence-activated cell sorting (FACS). To prevent RNA degradation, all solutions and reaction buffers except for ACME contained 1000 U/mL Recombinant RNase Inhibitor (Takara); all equipment coming in direct contact with the cell suspension was coated with the resuspension solution containing 40 U/mL of the inhibitor. For the same reason, all steps except for the initial dissociation and the EdU Click-iT™ reaction were performed on ice. Centrifugation was performed at 2000 × g and 4°C (Centrifuge 5910 R; Eppendorf).

#### Fluorescence-activated cell sorting (FACS)

EdU-treated cells and the corresponding control were sorted using BD FACSMelody™ Cell Sorter and FACSChorus v. 3.0 software (BD Biosciences). All other samples were processed using BD FACSAria™ III Cell Sorter and FACSDiva software v. 9.0.1 (BD Biosciences). In order to prevent RNase contamination, the devices were cleaned prior to cell sorting and maintained at 4°C. Sorting was performed in single-cell mode, using a 70 µm nozzle. Gates were set as shown in **Auxiliary Fig. 1-4** and 15000 cells from the EdU^+^ G1 and G2 gate (for the EdU-treated sample) and G1 and G2 gate (for the transcriptomic atlas) and 20000 cells from the P1 gate per sample (for the WT and *c-ops1^Δ8/Δ8^* dataset) were sorted into wells of a 96-well plate (MicroAmp Fast Optical 96-Well Reaction Plate with Barcode (0.1 mL); Life Technologies), containing reagents of the 10x Chromium single cell 3′ reagents kit v. 3.1 (10x Genomics): 18.8 µL RT Reagent B, 2.4 µL Template Switch Oligo and 2 µL Reducing Agent B; following the sorting, enough RT Enzyme C was added to each well to result in 52 µL of total volume.

#### Library preparation and sequencing

Libraries and sequences were generated at the Vienna BioCenter Next Generation Sequencing Facility. Suspensions were loaded onto a Chromium Controller chip (10x Genomics) and 3′ GEM libraries were constructed according to the manufacturer’s protocol. Library quality control was performed and the samples were sequenced and demultiplexed; sequencing was conducted on the NovaSeq 6000 sequencing system (Illumina), using the NovaSeq 6000 S4 Reagent Kit v. 1.5 (300 cycles) and following the XP workflow according to the manufacturer’s instructions.

#### General scRNA-seq analysis pipeline

Raw sequencing data (available at Dryad: https://datadryad.org/stash/share/x6pSympEtsaa7bUUflE4s3T1gxDZpy41JK3zxsTjtSU) were processed and mapped against version 1.0 of the *Platynereis* genome (*77*), following the default pipeline of CellRanger v. 7.0.1 or 8.0.0 (for the WT and *c-ops1^Δ8/Δ8^* dataset), forcing the program to recover 10000 cells from each sample. Mapping results and statistics are listed in **Auxiliary Fig. 5-8**. Unless otherwise stated, all further analyses were conducted using the Seurat package v. 4.4.0 (*33*), run on R software v. 4.4.1 through RStudio interface v. 2023.09. Comprehensive code with step-by-step headings can be found in the Dryad repository. In brief, barcodes, features and count matrices were converted into Seurat objects and quality control against multiplets and empty cells was performed for each library individually, removing cells with outlying read and feature numbers. Libraries were randomly downsampled to match the size of the library with the lowest number of cells, in order to ensure a faithful representation of cell type ratios across the maturation stages in the general and the subsequently subclustered datasets. As a control, the libraries in their original size were processed in the same manner as the downsampled ones (**fig. S12A, B**). Downsampled and control datasets were generated by merging the respective libraries. Merged datasets were normalized, the 2000 most variably expressed genes were identified for each library individually, the lists were collated and used to scale and center the data. For comparison, a downsampled and control dataset were generated following the standard Seurat integration pipeline: individual libraries were normalized, the 2000 most variably expressed genes were selected for each library, 2000 representative integration genes were selected for all libraries together and integration anchor cells were identified based on the gene list. Libraries were integrated and the resulting dataset was scaled. Principal component analysis (PCA) and UMAP reduction (*78*)were conducted on all datasets, discarding all but the first 40 or 35 principal components for merged and integrated datasets, respectively. Datasets were clustered and a clustering tree was generated, in order to select a resolution producing a stable number of clusters, using the Clustree package v. 0.5.0 (**fig. S12 C, D**) (*79*). For subcluster analyses, cell populations of interest were selected in each library separately and Seurat objects were generated from the subsets. The new libraries were then merged and the resulting dataset processed, annotated, analyzed and visualized in the same manner as the original dataset, with the exception that the list of 2000 most variable genes used for scaling was imported from the full subset, as opposed to from each individual library.

#### Transfer of cluster identities between datasets of the same origin

In **Fig. 1B, C**, cells belonging to clusters defined in the standard-integrated brain atlas dataset were highlighted on the UMAP reduction of the merge-integrated brain atlas dataset; this was achieved by selecting the cell populations of interest in the standard-integrated dataset and assigning them a unique color in the merge-integrated dataset.

#### Predictive dimensional reduction and annotation of new datasets

To project the EdU library, as well as the WT and *c-ops1^Δ8/Δ8^* dataset and its corresponding eye- and transcriptomically related cell subset (queries) onto the UMAP-reduced space of the existing brain atlas and eye- and transcriptomically related cell subeset (references), as well as predict the annotation of the cell populations in the new datasets, a Seurat object was generated out of the new libraries as previously described. Following this, a protocol based on the Seurat label transfer pipeline was followed: the UMAP model of the reference library was returned and the transfer anchors generated setting the reduction model to UMAP. For joint visualization of the reference and query, reference UMAP embeddings and predicted UMAP embeddings were extracted, collated, imported into a new dimensionally reduced object and visualized. Transfer anchors were then used for predicting the cell labels in the query (set to overarching cluster identities) and the predicted annotations imported into the object’s metadata. As a control, the *c-ops1^Δ8/Δ8^* and WT dataset and its eye- and transcriptomically related cell subset were dimensionally reduced and the projected cell labels were transferred following the standard pipeline described above.

#### Annotation of the adult *Platynereis* brain atlas

For annotation, we made use of a systematic comparison of genome-predicted transcripts against available sequence resources (*80*). Where available, entries were named based on BLASTN hits among published *Platynereis* sequences for which phylogenetic analyses had been performed; alternatively, best BLASTX hits in RefSeq proteins of selected landmark organisms was used, adding the suffix -like to the name if no phylogenetic analysis was performed. Entries with no BLAST hits were called by their locus identifier (XLOC number). For dataset annotations, marker genes of each cluster were computed using the nonparametric Wilcoxon rank sum test (default setting) and gene co-expression analysis was run using the fcoex package v. 1.10.0 (*81*) (**fig. S1B-N, Auxiliary file Tab 1, Auxiliary file Tab 2**). An fcoex object was created from the normalized count matrix and Seurat cluster identities; gene co-expression modules and correlation networks were generated and the dataset reclustered in order to visualize the module-based populations. The results of cluster marker identification and gene co-expression analysis were cross-compared with literature on *Platynereis* and vertebrate cell types and known gene expression domains; this was used to assign putative identities to clusters or group clusters into an overarching identity.

#### Differential expression analysis

In order to identify the genes most differentially expressed between EdU-treated and untreated worms (**Auxiliary file Tab 4**), as well as between non-reproductive and reproductive worms (**Auxiliary file Tab 7**), differential expression was performed between the corresponding libraries. Merged datasets were generated containing the libraries of interest, as described in “General scRNA-seq analysis pipeline”. Cells of respective libraries were combined into two lists, which were used to generate new identities in the integrated datasets. The identities were then treated as two clusters and the genes most differentially expressed between them were computed using the default parameters. To determine the genes differentially expressed between EPs and all brain cells, as well as support cells and all brain cells in **fig. S4G, H** and **Auxiliary File Tab 5**, the procedure was followed as described, with the exception that the negative binomial distribution statistical test was used, in line with methods used in ref. (*38*).

#### Pseudotime analysis

To determine and visualize lineage trajectories, selected cell populations were analyzed in pseudotime. The Seurat object was manually converted into a Monocle3 object and the differentiation trajectory was predicted following the standard vignette of the package Monocle3 v. 1.2.9 (*43*); cells were then ordered from least to most differentiated along the trajectory, selecting the Monocle3-suggested root nodes with the expression of expected marker genes as the starting points of each individual leaf of the trajectory. Marker genes were selected either for their known involvement in neurogenesis and proliferation in *Platynereis* or through differential expression analysis between EdU and premature or sampling time-matched control library in the eye- and transcriptomically related cell subset (**Auxiliary file Tab 4**). Expression of differentially expressed genes was individually screened regardless of putative gene function and very broadly and very sparsely expressed genes were excluded; the resulting genes were plotted as a cumulative module to indicate populations derived from proliferating cells.

#### Transcriptomic data visualization

For visualization of gene expression, non-biological zero values were imputed using the ALRA package v. 1.0.0 (*82*), unless otherwise stated. Color palettes used in plotting were selected from the package Paletteer v. 1.5.0. All plots were exported in the PDF format. Gene expression was plotted either individually or as an average expression level of several genes generated using the standard function for calculating cumulative module scores in Seurat. Dotplots were generated with the help of scCustomize package v. 1.1.1; otherwise, the standard plotting functions of respective packages were used with the following exceptions: in **Fig. 1C**, cluster identities transferred from the integrated dataset onto the merged dataset were plotted and exported individually, then overlaid in Adobe Photoshop 2024 as separate layers, using the layer blending option Linear Dodge (Add). Bubble plots in **Fig. 1L** and **fig. S2C** were generated in Microsoft Excel (Microsoft Office Professional Plus 2019), from differential expression analysis results exported as comma separated files. Stacked bar plots were generated using Microsoft Excel (Microsoft Office Professional Plus 2019). In **Fig. 2M** and **fig. 6D**, the differentiation trajectory and pseudotime-ordered cells were plotted and exported separately, then overlaid in Adobe Illustrator 2023. In **Fig. 3C, D**, cells colored in UMAP-reduced space were additionally outlined in Adobe Illustrator 2023. Scatterplots in **Fig. 4G-K** were generated from unimputed gene expression values that were exported as comma-separated files for each library individually and plotted in GraphPad Prism software v. 10.1.0 as superimposed scatter plots with manually added significance values calculated as described in “Differential expression analysis”. The order of cells in all scatterplots in **Fig. 4** was randomly shuffled in Adobe Illustrator 2023, to ensure faithful representation of all datasets in a plot. Where applicable, genome identifier numbers were substituted with gene names in Adobe Illustrator 2023

#### Worm eye size scoring and statistical analysis

To estimate the eye size in reproductive animals, female and male worms were anesthetized as previously described and their heads imaged with the SMZ18 stereomicroscope, DS-Ri2 camera and NIS Elements imaging software (Nikon) under the bright field setting. For each individual, head area and the left and right eye areas were calculated individually in a Z-projected image using Fiji v. 2.9.0 (*83*) by manually selecting the region of interest. Eye areas were then summed up. Head area was determined following the head border, excluding the palpae, cirri and the area posterior to the nuchal organ commissure. Total eye area was normalized to the head area for each worm. Data were plotted and statistically analyzed in GraphPad Prism software v. 9.5.1 with an unpaired t test. To compare the eye size between WTs and *c-ops1^Δ8/Δ8^* animals, the worms were processed as described in “EdU labeling” and “EdU detection, HCR and immunohistochemistry (IHC)”; the nuclear stain channel was imaged as described in “Microscopy and image processing” and the eye areas calculated in a Z-projected image in Fiji v. 2.9.0 following the borders of both posterior eyes. Data points were plotted as replicates (two posterior eyes per worm) and analyzed with a nested t-test in GraphPad Prism software v. 10.1.0; all measurements available in **Auxiliary file Tab 3**.

#### EdU labeling

Proliferating cells were labeled using an EdU incorporation assay as previously described (*57*). Briefly, selected worms were transferred into small glass beakers and incubated with 10 µM EdU dissolved in worm culture water for 4, 16 or 96 hours; in the case of a 16-hour pulse, worms were sampled at ZT 10 and incubated overnight, otherwise as stated in “Worm staging and sampling”. For experiments involving a chase period, EdU solution was discarded following the completion of a pulse, the animals and the beaker were rinsed several times with worm culture water and 100 µM thymidine dissolved in worm culture water was added. The animals were left to incubate for 168 hours (1 week). Control worms were incubated in pure worm culture water during the pulse and thymidine solution during the chase.

#### EdU detection, HCR and immunohistochemistry (IHC)

Depending on the experiment, EdU detection was either conducted on single-cell suspensions for scRNA-seq experiments or on whole worm heads for imaging experiments; for the latter, EdU was either detected on its own following the manufacturer’s protocol, or in conjunction with the protocol for simultaneous HCR, IHC and EdU (*57*). In short, following the pulse and the optional chase, head tissue was sampled, fixed and dehydrated. The following day, samples were left to incubate overnight in a solution containing HCR probes against transcripts of interest; next, HCR amplifiers were added and optionally, EdU Click-iT™ reaction was performed according to the manufacturer’s protocol. In case that IHC was being conducted in parallel, primary antibody against the protein of interest was added to the reaction mix and left to incubate overnight. The following day, HCR amplification was terminated and optionally, the secondary antibody was added; after an overnight incubation, the samples were washed, mounted and imaged. Control samples received HCR probes against unrelated genes designed for non-annelid species and no primary antibody.

#### EdU signal quantification

To quantify proliferating cell numbers, worms were labeled with either a 16-hour (for comparison between stages) or 4-hour EdU pulse (for comparison of eye and posterior oval shaped-domain signal between *c-opsin1* mutants and WTs). The former experiment was imaged with the Axio Imager Z2 upright microscope (Zeiss), using the Plan-Neofluar 10x/0.3 (dry) lens, AxioCam MRc5 camera and ZEN Blue software v. 3.3. The latter experiment was imaged as described in “Microscopy and image processing”. In Fiji v. 2.9.0 (*83*), all images were processed as described in “Microscopy and image processing”. Regions of interest were selected in Z-projected images as described in “Worm eye size scoring and statistical analysis”. EdU-labeled nuclei within the region of interest were manually counted, and in the experiment comparing EdU signal between maturation stages, the number was normalized to the head area. Data points were plotted individually or as replicates (two posterior eyes per worm) and statistical significance was calculated in GraphPad Prism software v. 10.1.0 with a one-way ANOVA test and Tukey’s multiple comparisons correction or a nested t-test, for the comparison of EdU signal between maturation stages or between *c-opsin1* mutants and WTs, respectively; all measurements available in **Auxiliary file Tab 3**.

#### HCR probes and antibodies

All HCR probes against genes of interest (**Auxiliary file Tab 10**) were designed using the HCR Probe Maker software (insitu_probe_generator v. 0.3.2), installed with Python v. 3.10.2 and JupyterLab v. 3.4.0 (*84*), with the exception of *Pladu_prox*, which was designed and manufactured by Molecular Instruments. Using CLC Main Workbench v. 22, probe pairs were mapped against the sequences of interest and those binding to highly polymorphic regions were discarded. B1 amplifier coupled to Alexa Fluor™ 546 fluorophore and B2 amplifier coupled to Alexa Fluor™ 647 fluorophore were used for imaging.

#### Microscopy and image processing

Visualization and imaging of fluorescently labeled worm heads were achieved with the LSM 700 inverted confocal microscope (Zeiss), using EC Plan-Neofluar 10x/0.3 (dry), Plan-Apochromat 20x/0.8 (dry), LD LCI Plan-Apochromat 25x/0.8 (oil immersion), Plan-Apochromat 40x/1.3 (oil immersion) and Plan-Apochromat 63x/1.4 (oil immersion) lenses and the ZEN (black edition) desk software v. 2.3. In Fig. 3, the imaging of unprocessed worm heads for size comparison was achieved with the Axio Imager Z2 upright microscope (Zeiss), using the Plan-Neofluar 10x/0.3 (dry) lens, AxioCam MRc5 camera and ZEN Blue software v. 3.3, set in brightfield mode. Parameters for acquisition were set according to a representative treated sample and applied to other samples belonging to the same experimental batch. Images were processed with Fiji v. 2.9.0 (*83*), optimizing brightness and contrast for each channel separately, generating a Z-projection out of selected slices in an image stack using the Maximal Intensity option and optionally enhancing the sharpness of the resulting image. For some image stacks consisting of a larger number of slices, the Stack Contrast Adjustment plugin was used prior to Z-projecting the image, in order to compensate for the loss of fluorescence intensity in deeper tissue layers due to photon scattering (*85*). Settings were adjusted to best fit a representative treated sample and applied to the control sample of the same batch; for some of the samples not used for quantification, the settings of the nuclear stain channel were adjusted individually due to the differences in tissue thickness and photon scattering at short wavelengths. Outlines were added and panels were arranged in Adobe Illustrator 2023.

#### Gene ontology term enrichment analysis

GO terms linked to genes differentially expressed between non-reproductive and reproductive worms were obtained using the web server ShinyGO v. 080 (*86*). *Platynereis* genes were matched against the human proteome and where applicable, *Homo sapiens* UniProt IDs were used for comparison, as described (*80*). Hits of all genes were set as the background, and the hits of differentially expressed genes as the query; the search was filtered for biological processes and the analysis performed using the default settings of the web server. Highest ranking terms were visualized with the built-in chart function. Conversely, to probe for enrichment of terms related to molecular mechanisms and cellular processes of interest, a reverse search was performed in the AmiGO 2 Ontology database (*87*) using default filters, and resulting GO terms affiliated with or containing the keywords were looked up in the results of the enrichment analysis (**Auxiliary file Tab 8, 9**).

#### Phylogenetic tree construction

To determine potential orthologs of vertebrate and *Drosophila* protein coding sequences in the worm, a cDNA database with genome-predicted transcripts was used. For the reported phylogenies, reference sequences were re-assembled from available cDNA resources of the investigated laboratory strains (*28*, *38*, *88*), also correcting cases of missing or mis-predicted genes (*tob* and *lamin*). Best reciprocal matches from *Mus musculus* and *Drosophila melanogaster* were retrieved from UniProt, using blastx/tblastn queries in CLC Main Workbench v. 22; transcript variants with the longest reading frame were translated to protein, and additional protein hits from UniProtKB/SwissProt and RefSeq were obtained to identify possible matches in additional reference species (*Homo sapiens, Mus musculus, Gallus gallus, Xenopus laevis, Xenopus tropicalis, Strongylocentrotus purpuratus, Saccoglossus kowalevskii, Capitella teleta, Crassostrea gigas, Mizuhopecten yessoensis, Pomacea canaliculata, Biomphalaria glabrata, Drosophila melanogaster, Tribolium castaneum, Apis mellifera, Daphnia pulex, Priapulus caudatus, Lottia gigantea* and *Helobdella robusta*) Hit sequences were aligned in CLC Main Workbench v. 22 and exported in FASTA format for phylogenetic tree construction. Maximum likelihood analysis was run on the sequences using the IQ Tree web server and the best substitution model was selected by the web application W-IQ-TREE, using the ModelFinder algorithm operating in jModelTest mode (*89*, *90*). A stochastic algorithm was implemented to achieve a high likelihood ratio in order to locate local optima in the tree space, and a bootstrap approximation was performed with UFBoot2 (*91*), using the default settings and performing 1000 iterations. Trees were visualized and arranged using the Interactive Tree of Life tool (*92*); trees were manually rooted using a protein outgroup obtained using the second-best hit in the mouse, *Drosophila* or *Platynereis* proteome as a starting point. Tree construction was reiterated several times in order to eliminate duplicates and outliers, resulting in a stable tree containing sequences of all relevant reference groups. A comprehensive list of proteins used in the final phylogenetic tree assembly and their respective accession numbers in different reference organisms can be found in **Auxiliary file Tab 11**. Sequences of *Platynereis* genes identified/validated in this manner are available under the following GenBank identifiers: XXX – foxo, XXX – btg1/2, XXX – btg3/4, XXX – id/emc, XXX – nf, XXX – ki67, XXX – fut10.

#### Protein domain analysis

Phylogenetic trees were supported with comparative analyses of functional domains in orthologous proteins from selected organisms. Simple Modular Architecture Research Tool (*93*) was used to identify and visualize domains in the sequences of interest, which were subsequently annotated by retrieving family and domain information for each individual protein from the UniProt database of the respective organism.

## Acknowledgments

We are grateful to Andrij Belokurov, Margaryta Borysova and Netsanet Getachew for routine worm cultures and genotyping support, Lena Stumbauer for practical help, as well as all members of the Tessmar-Raible and Raible labs for constructive discussions.

## Funding

This work was supported by

Helmholtz Society, distinguished professorship by the Alfred Wegener Institute Helmholtz Centre for Polar and Marine Research (K.T-R)

H2020 European Research Council, ERC Grant Agreement #819952 (K.T-R)

Austrian Science Funds (FWF), SFB F78 (F.R., K.T-R)

University of Vienna Research Platform SinCeReSt (F.R.)

None of the funding bodies was involved in the design of the study, the collection, analysis, and interpretation of data or in writing the manuscript.

## Author contributions

Conceptualization: NM, FR, KTR

Methodology: NM, CLVG, GA, FR, KTR

Investigation: NM, CLVG, GA, FR, KTR

Visualization: NM, CLVG, GA

Funding acquisition: FR, KTR

Project administration: FR, KTR

Supervision: FR, KTR

Writing – original draft: NM, FR, KTR

Writing – review & editing: NM, CLVG, GA, FR, KTR

## Data and materials availability

All data are available in the main text, the supplementary material, and auxiliary figures/files and data deposited at the Dryad repository: https://datadryad.org/stash/share/x6pSympEtsaa7bUUflE4s3T1gxDZpy41JK3zxsTjtSU

## Supplementary Materials

**Other Supplementary Materials for this manuscript include the following:**

Dryad archive: https://datadryad.org/stash/share/x6pSympEtsaa7bUUflE4s3T1gxDZpy41JK3zxsTjtSU

Containing:

Auxiliary file (Tab 1 to 11)

Auxiliary Fig. 1 to 8

R code

R objects

Raw scRNA-seq data

**Fig. S1.**
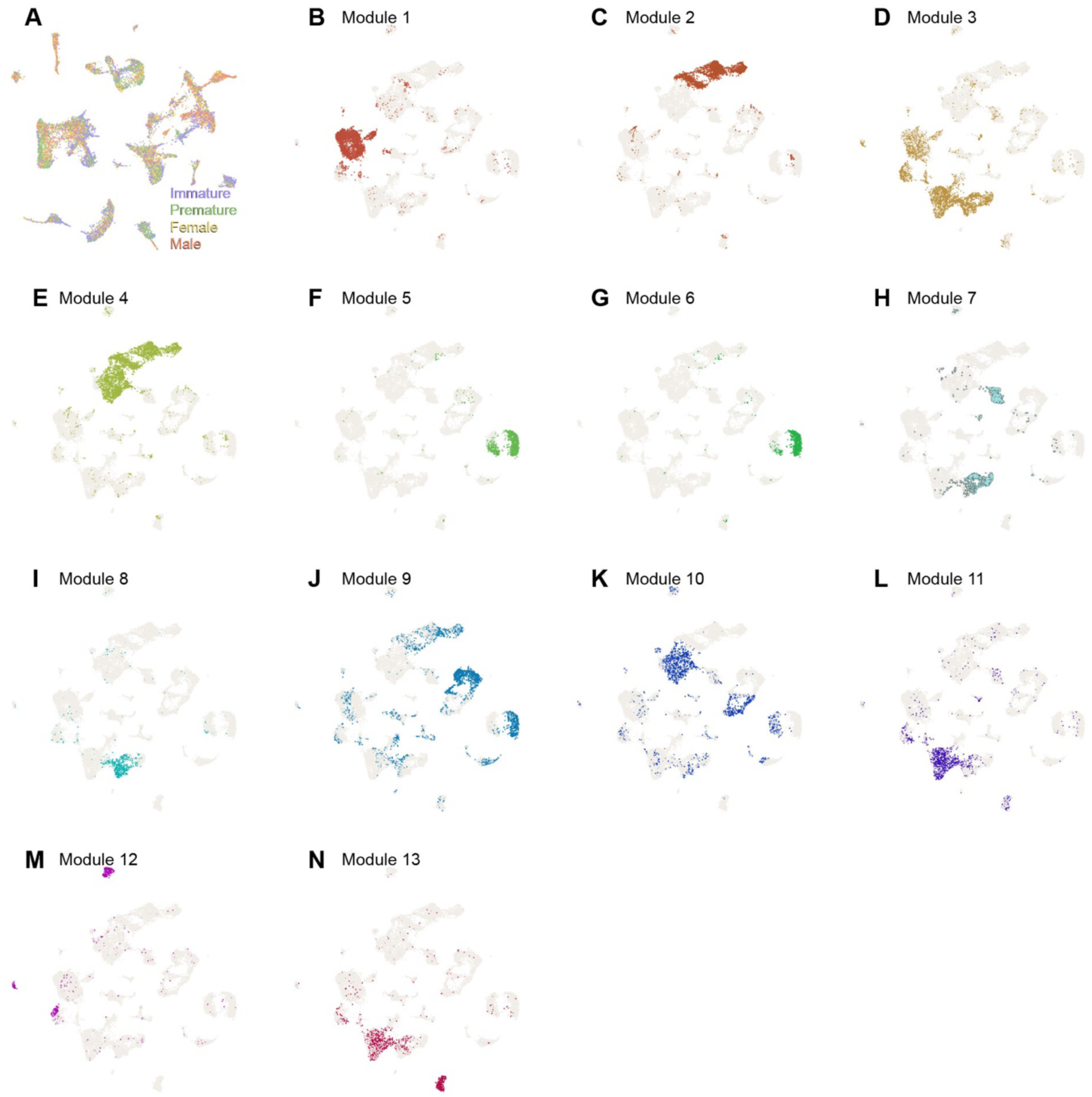
Integration and annotation of the brain atlas. (**A**) Integration of immature, premature, female and male libraries performed following the standard pipeline of the R package Seurat. (**B-N**) Independent identification of gene co-expression modules used for manual overarching cluster identity assignment.

**Fig S2.**
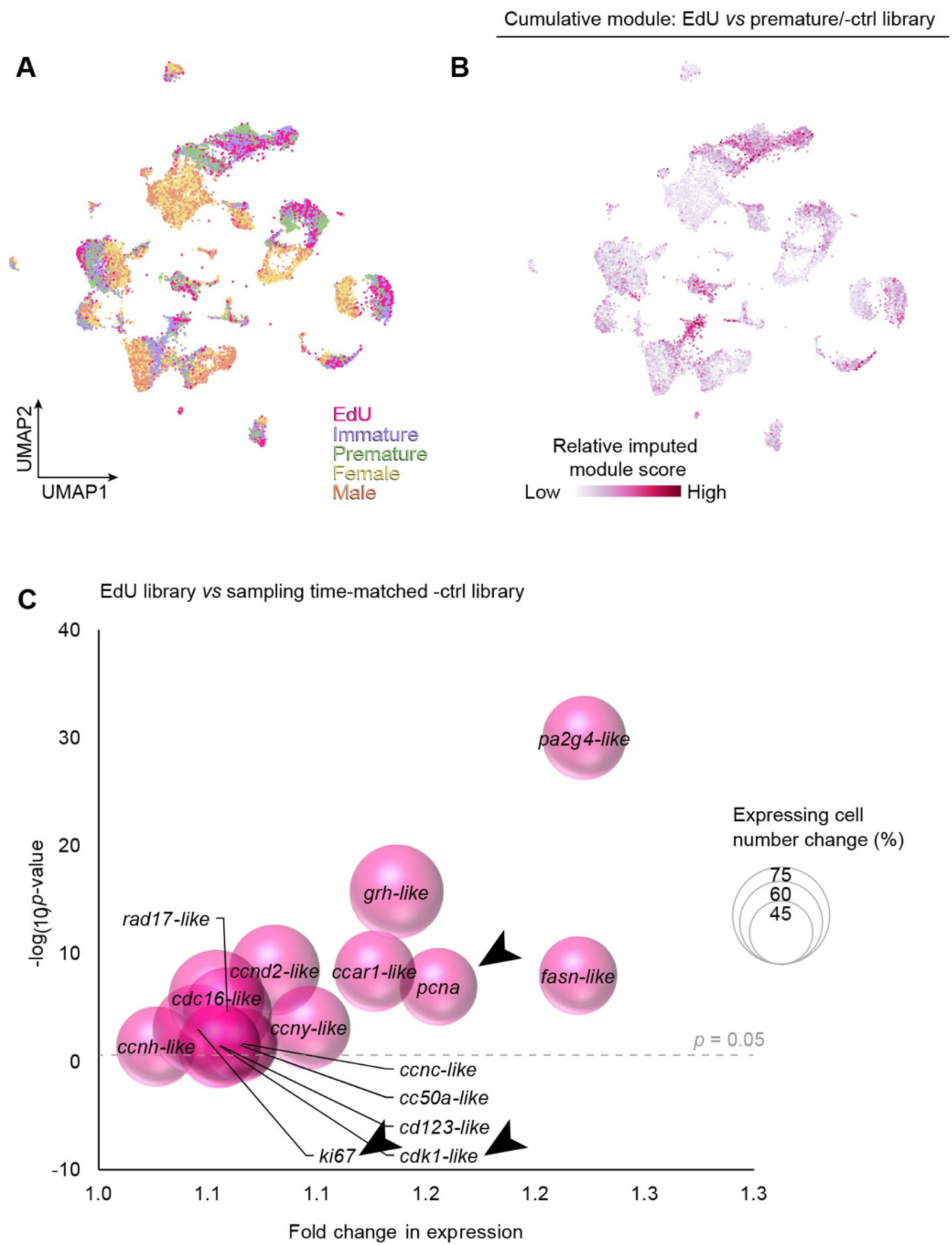
Determination of genes enriched in the proliferating cell population. (**A**) EdU scRNA-seq library projected over UMAP-reduced space of the merge-integrated brain atlas dataset. (**B**) Cumulative module of *bona fide* neurogenesis- and proliferation-associated genes differentially expressed between EdU library and premature library (**Fig. 1L**) and EdU library and sampling time-matched control library in (**C**). Differentially expressed candidates were selected on the criterion of playing a role in cell cycle regulation and neurogenesis in humans and mice, as well as neurogenesis, neural commitment and cell proliferation in *Platynereis* and *Drosophila*; genes whose orthologs have not yet been investigated in *Platynereis* are presented as best reciprocal matches of mouse and human genes with the added suffix -like; see “Annotation of the adult *Platynereis* brain atlas” in Methods for more details. (**C**) *Bona fide* neurogenesis- and proliferation-associated genes differentially expressed between the EdU library and the sampling time-matched control library. Only genes surpassing the indicated *p*-value threshold are displayed. Arrowheads indicate genes mentioned in main text. Statistical significance: Wilcoxon Rank Sum test.

**Fig. S3.**
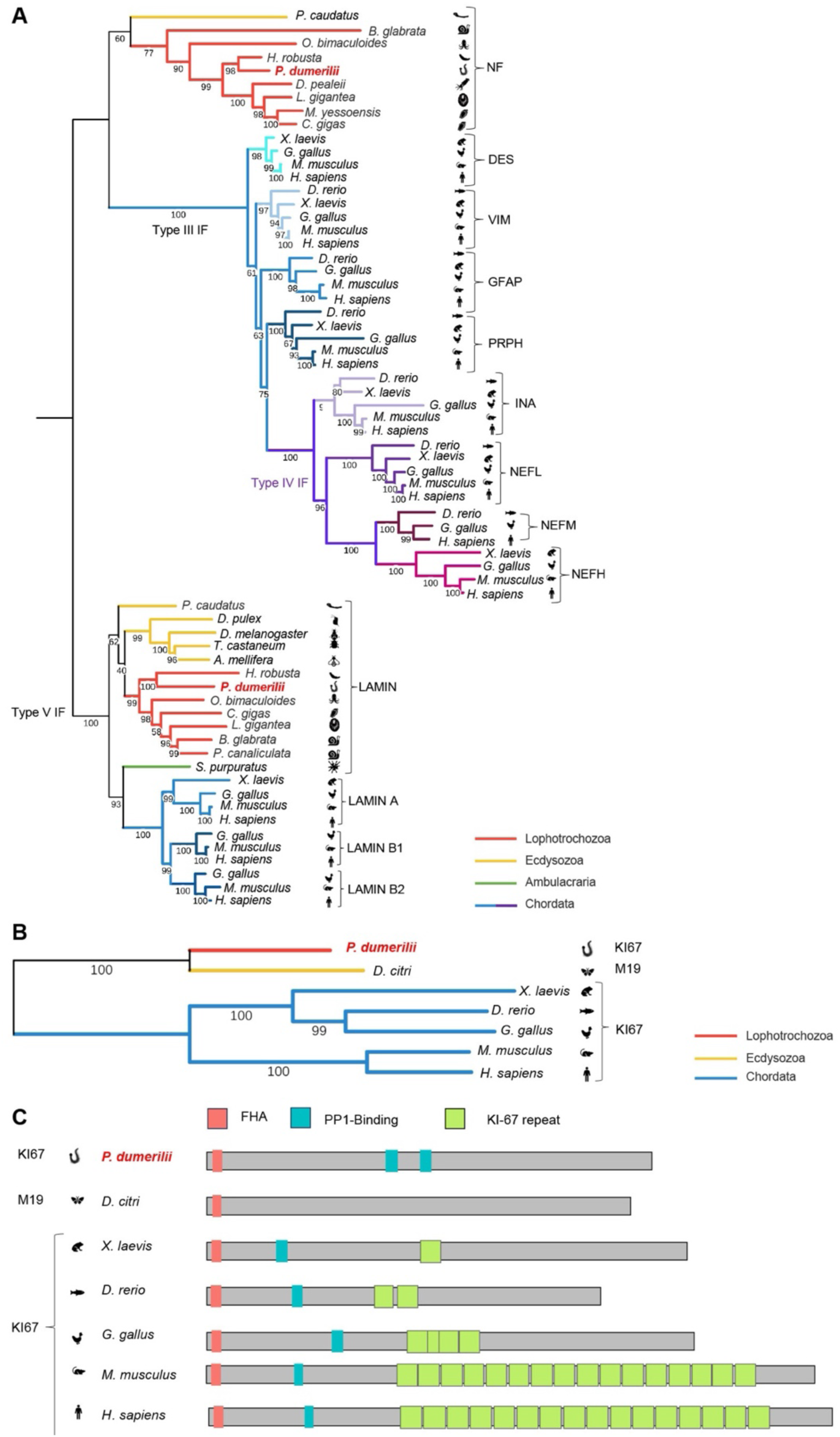
Phylogeny and expression of *bona fide* neurogenesis- and proliferation associated genes. (**A**) Phylogeny of NF, the newly identified *Platynereis* ortholog of GFAP; bootstrapping percentage shown in internal branches. Lamin proteins were used as an outgroup. (**B**) Phylogeny of Ki67; bootstrapping percentage shown in internal branches; unrooted tree. (**C**) Schematic representation of protein domains corresponding to proteins shown in (**B**).

**Fig. S4.**
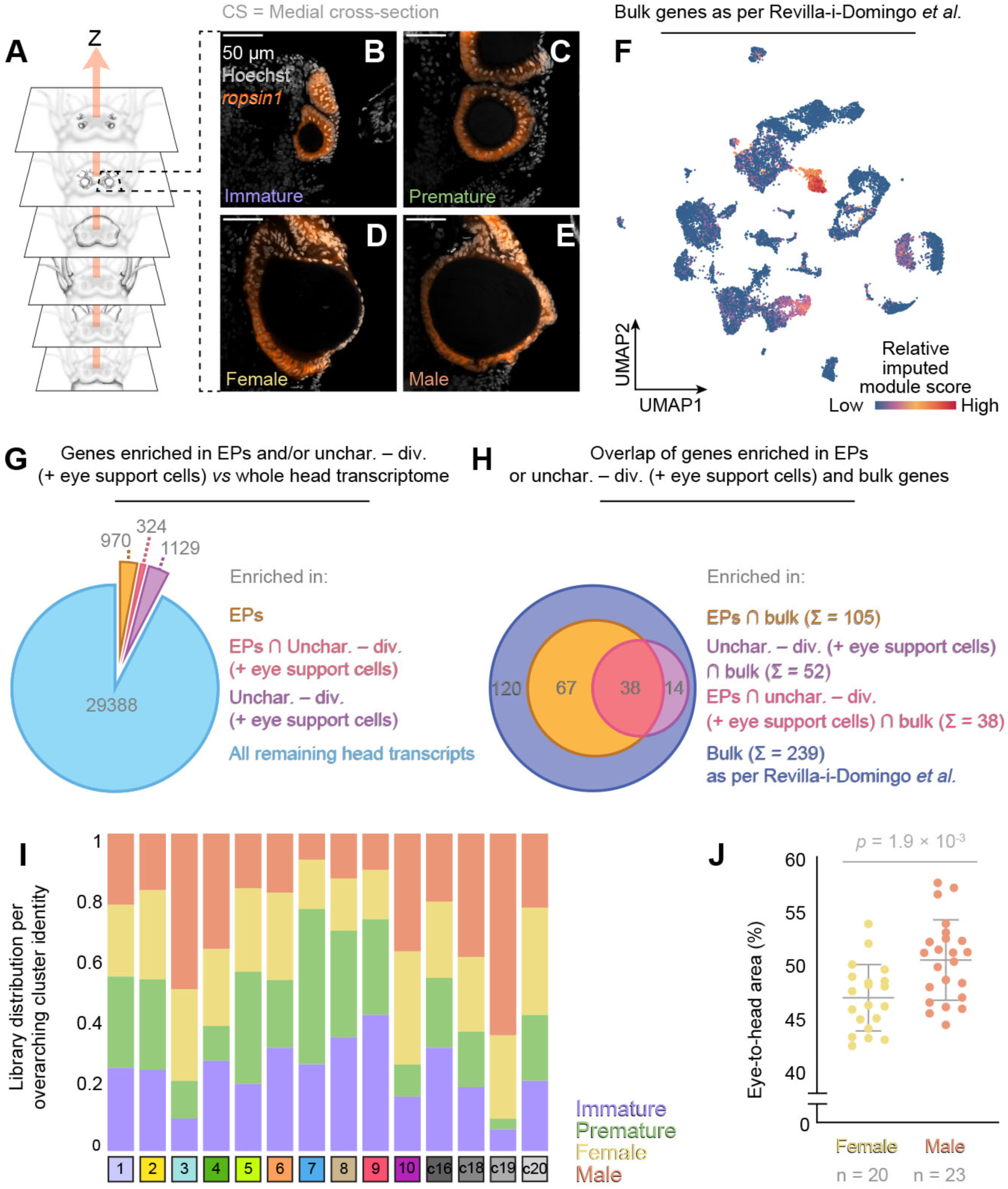
*In situ* and *in silico* identification of eye growth. (**A**) Schematic representation of the focal plane in (B-E). (**B-E**) *In situ* HCR staining of *r-opsin1* (red) in (**B**) immature, (**C**) premature, (**D**) female and (**E**) male eye; images are 1 µm thick. (**F**) Cumulative gene module of genes enriched exclusively in eye photoreceptors and jointly enriched in eye photoreceptors and trunk *r-opsin1* expressing cells, as detected by bulk RNA-seq in ref. (*1*); in further text referred to as “bulk”. Relative Venn diagrams of (**G**) genes enriched in EPs and uncharacterized – diverse (+ eye support cells) against the whole head transcriptome, as well as their overlap, and (**H**) the overlap of genes exclusively or jointly enriched in EPs and/or uncharacterized – diverse (+ eye support cells) and bulk genes. (**I**) Representation of immature, premature, female and male scRNA-seq libraries in overarching cluster identities of the brain atlas. (**J**) Comparison of eye size of reproductively mature female (n = 20) and male (n = 23) worms. Values are displayed as cumulative area of four eyes per animal normalized to the head area. Statistics: unpaired t-test.

**Fig. S5.**
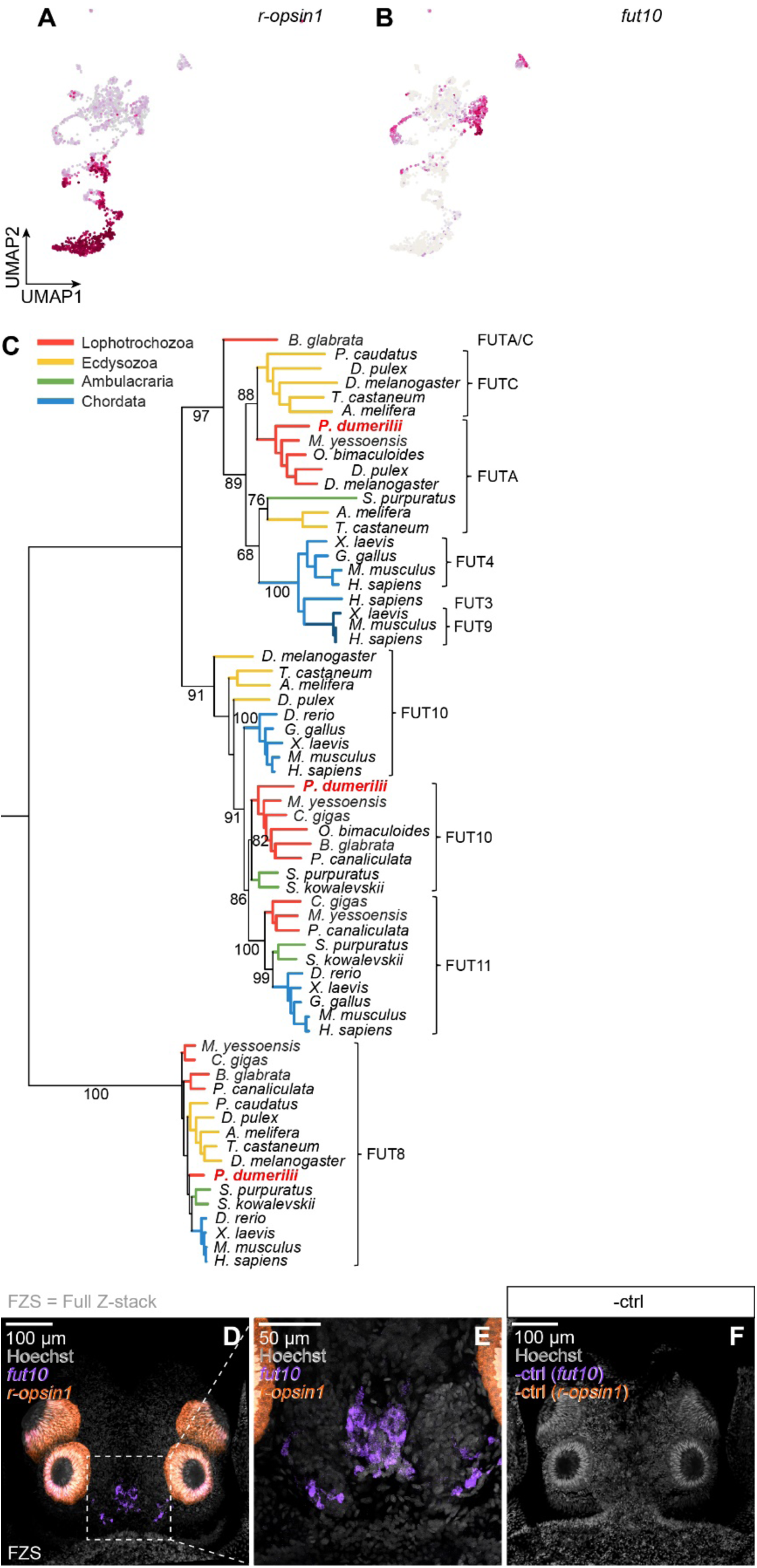
Phylogeny and expression of FUT10/*fut10*. Expression of (**A**) EP marker *r-opsin1* and (**B**) newly identified eye support cell marker *fut10*. (**C**) Phylogenetic tree of FUT10; bootstrapping percentage shown in internal branches. Tree was rooted using FUT8 as an outgroup. (**D-F**) HCR detection of *fut10* expression in the head and corresponding negative control. Microscopy images are of 51.44, 24.94 and 36.73 µm respective thickness.

**Fig. S6.**
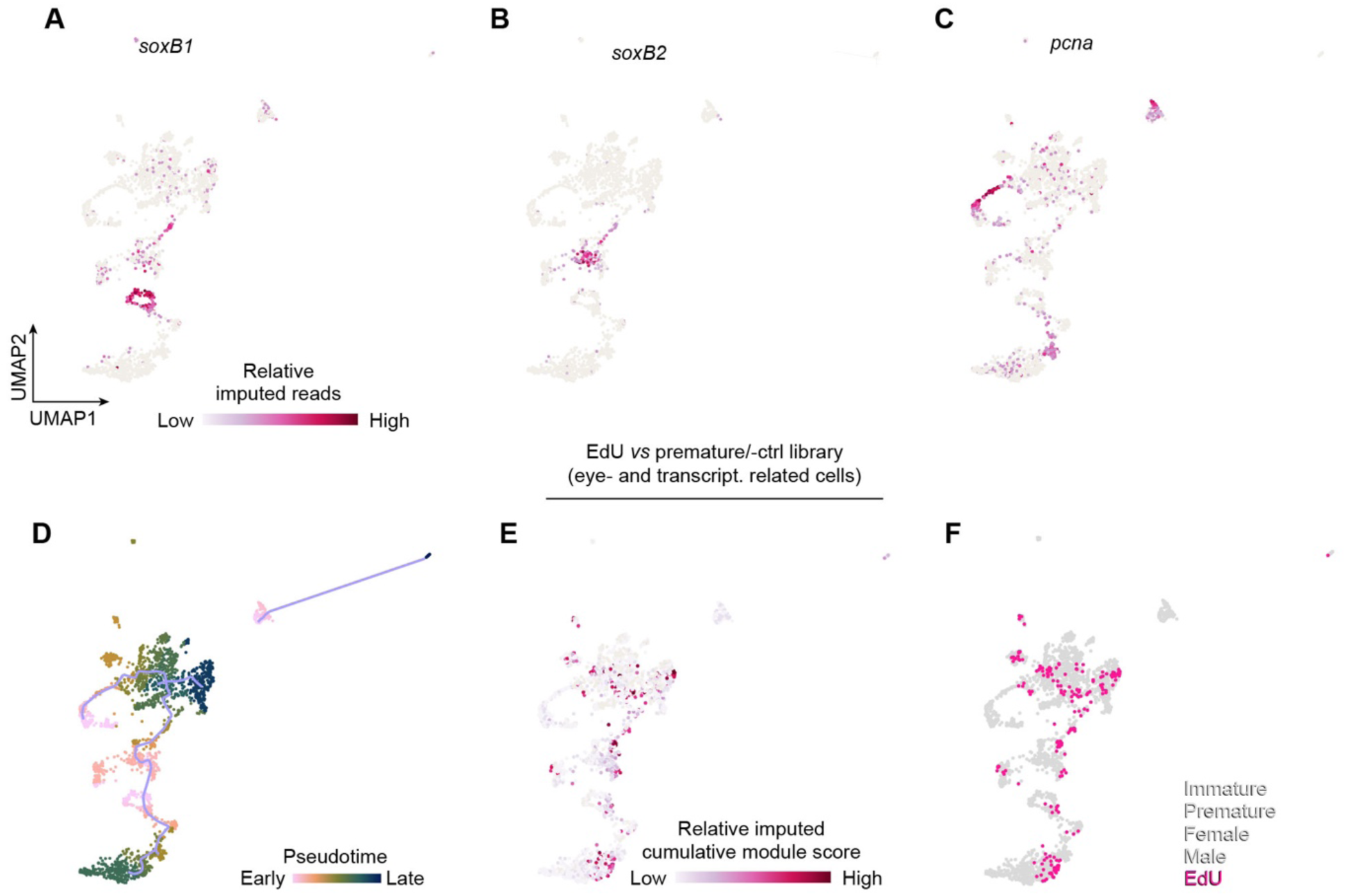
Likely source populations and differentiation trajectories of stem cells in the eye. **(A-C)** Expression of neurogenesis- and proliferation-associated markers (A) *soxB1*, (B) *soxB2* and (C) *pcna* in the eye- and transcriptomically related cell subset. (**D**) Putative differentiation and pseudotime ranking of cells corresponding to the whole eye- and transcriptomically related cell dataset; due to the transcriptomic similarities of all cells in the upper partition of the graph, the eye support cells appear as progeny of two trajectory root nodes. (**E, F**) Differentiation trajectories inferred by the EdU^+^ dataset; (E) Expression of a cumulative module computed from genes differentially expressed between the eye- and transcriptomically related cell subset of the EdU^+^ library and the premature and sampling time-matched control libraries, respectively (F) Visualisation of cell transcriptomes from the EdU^+^ library in a joint dataset consisting of the immature, premature, male, female and EdU^+^ transcriptomes. Note that due to a 96-hour pulse, most labeled cells are lineage-committed or differentiated progeny of stem cells.

**Fig. S7.**
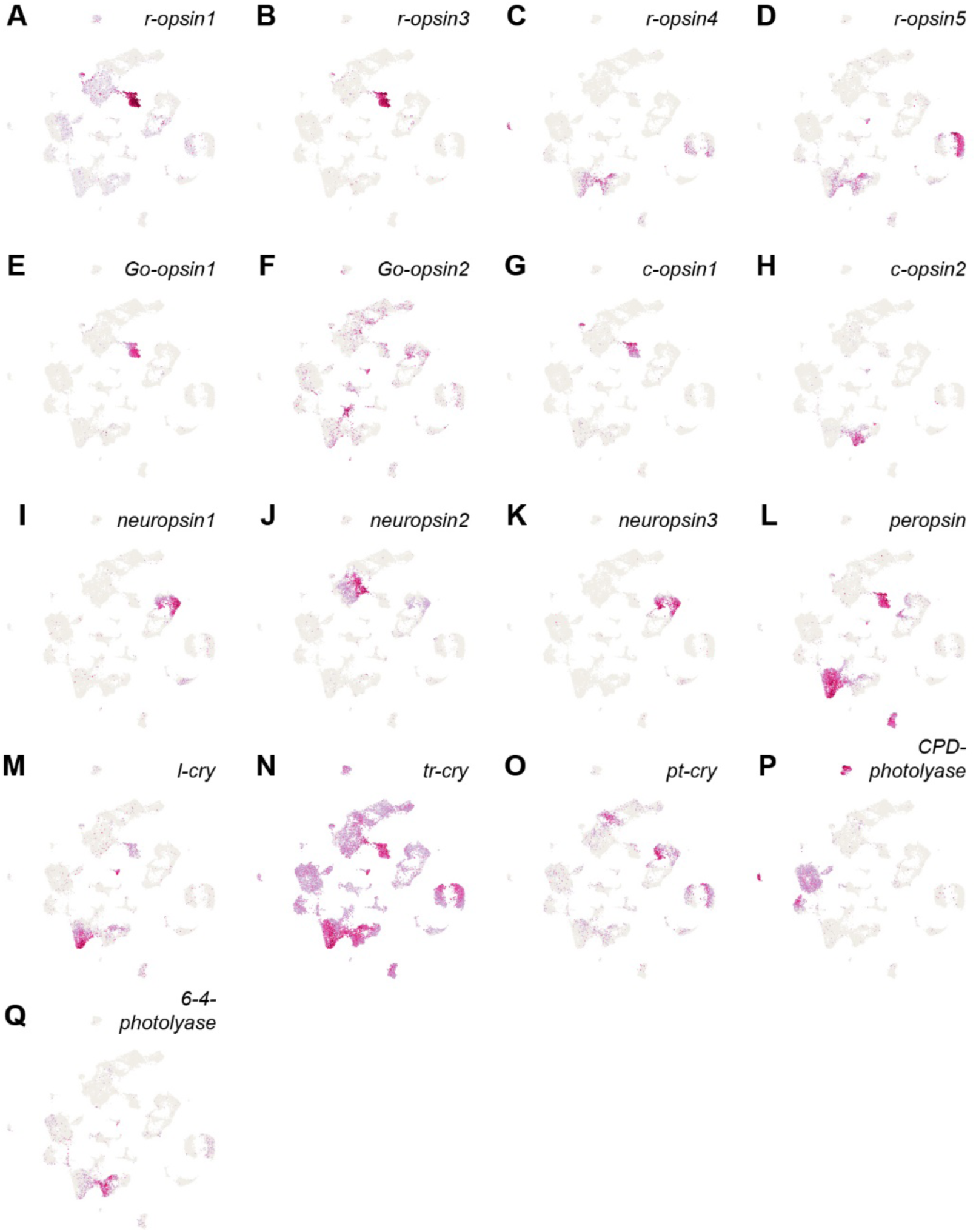
Expression of known photoreceptor genes in the *Platynereis* brain atlas. (**A-Q**) Expression predicted by scRNA-seq of currently known *Platynereis* photoreceptor genes in the brain atlas. *r-opsin2* and *cry-DASH* were omitted from the figure due to a lack of reads and the absence of an appropriate gene model in the current genome, respectively.

**Fig. S8.**
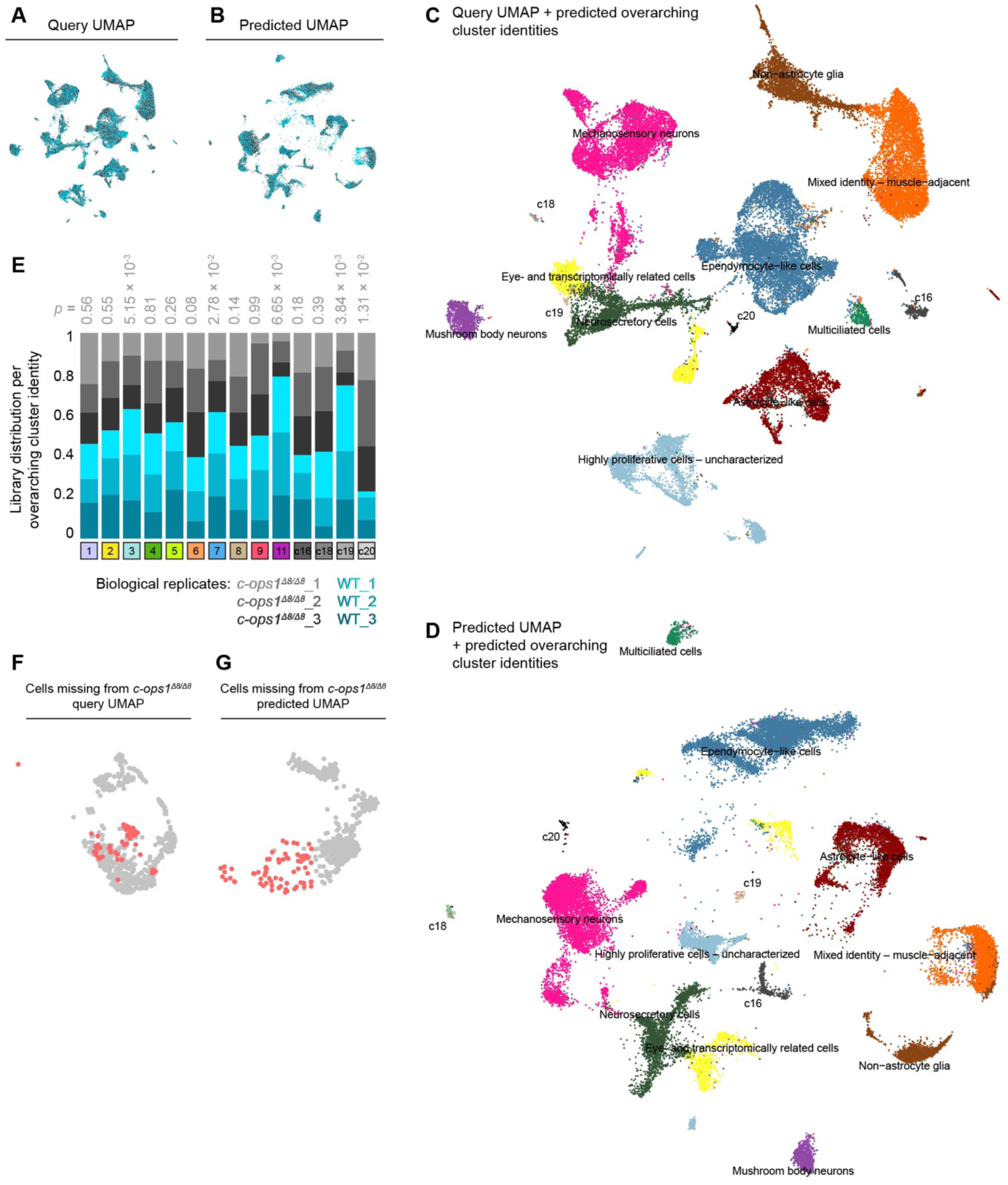
Integration and annotation of the *c-ops1^Δ8/Δ8^* and WT dataset. UMAP-projections of the *c-ops1^Δ8/Δ8^* and WT dataset (**A**) obtained through standard dimensional reduction of the dataset (query) and (**B**) projected over the brain atlas UMAP by predicting the UMAP-coordinates of the query using the coordinates of the brain atlas as a reference. Predicted overarching cluster identities on the (**C**) query UMAP-reduction and (**D**) predicted UMAP-reduction. (**E**) Representation of *c-ops1^Δ8/Δ8^* and WT libraries in predicted overarching cluster identities of the brain atlas. A subset of the *c-ops1^Δ8/Δ8^* and WT dataset corresponding to EPs and matching eye aNSCs/PCs visualized as a (**F**) query UMAP-reduction and (**G**) predicted UMAP-reduction using the UMAP-coordinates of the eye subset of the brain atlas as a reference. Highlighted cells are absent from *c-ops1^Δ8/Δ8^* libraries.

**Fig. S9.**
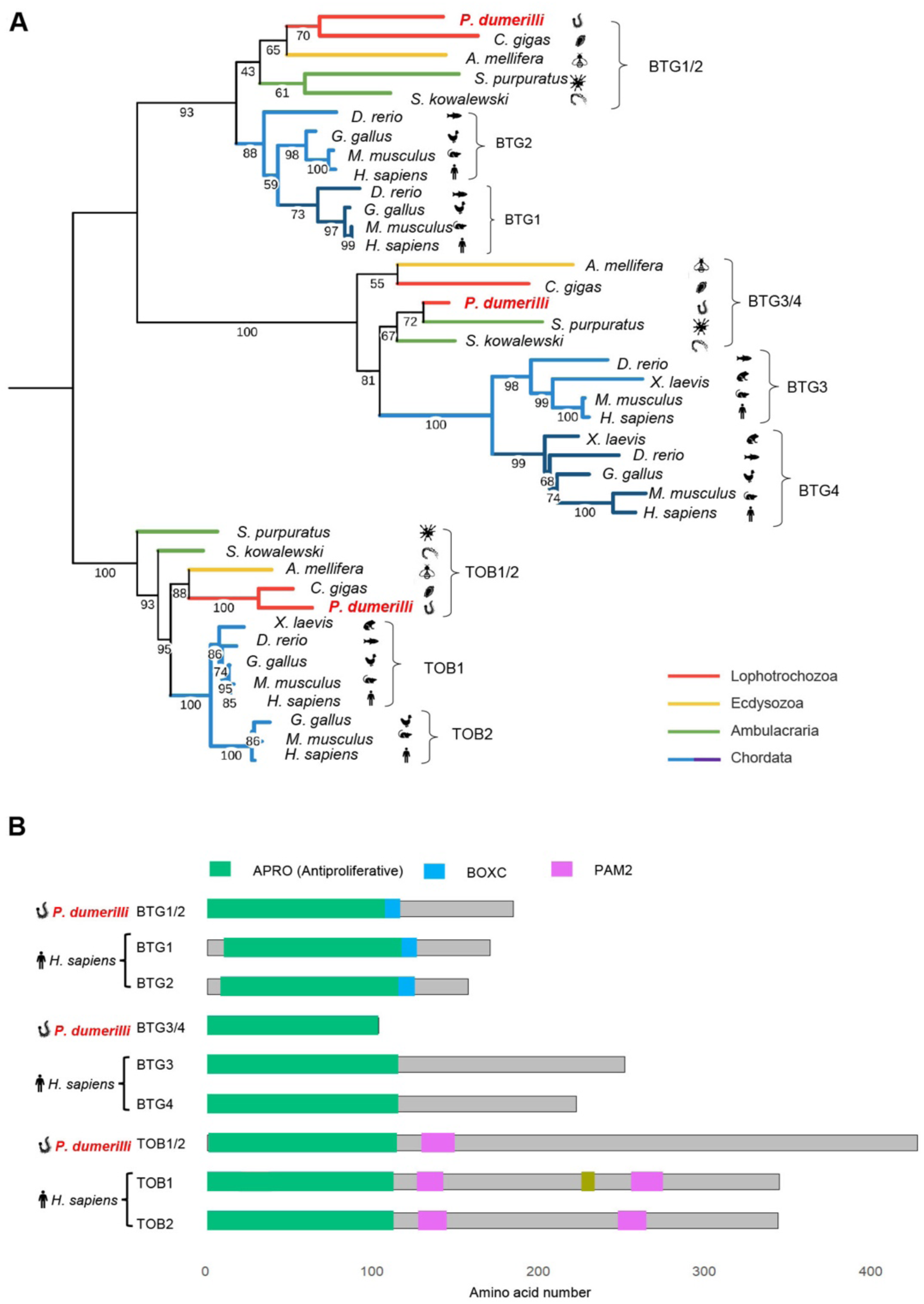
Phylogeny and domain architecture of BTG1/2 and BTG3/4. (**A**) Phylogenetic tree of the BTG/TOB protein family; statistical support by bootstrapping percentage shown in internal branches. Tree was rooted TOB family as an outgroup. (**B**) Schematic representation of protein domains corresponding to proteins shown in (**A**) in *P. dumerilii* and *H. sapiens*.

**Fig. S10.**
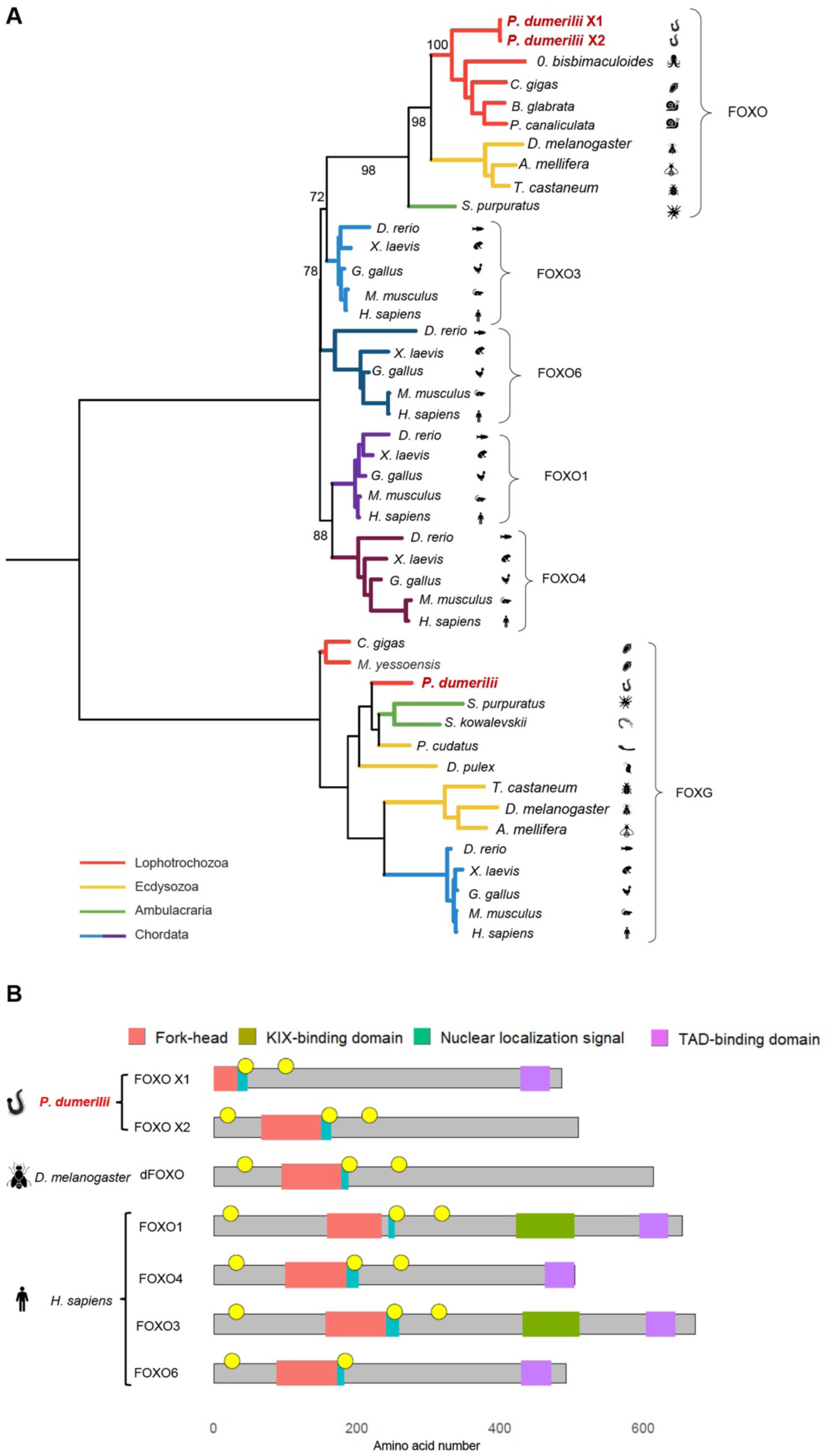
Phylogeny and domain architecture of FOXO. (**A**) Phylogenetic tree of FOXO; statistical support by bootstrapping percentage shown in internal branches. Tree was rooted FOXG as an outgroup. (**B**) Schematic representation of protein domains corresponding to proteins shown in (**A**) in *P. dumerilii*, *D. melanogaster* and *H. sapiens*.

**Fig. S11.**
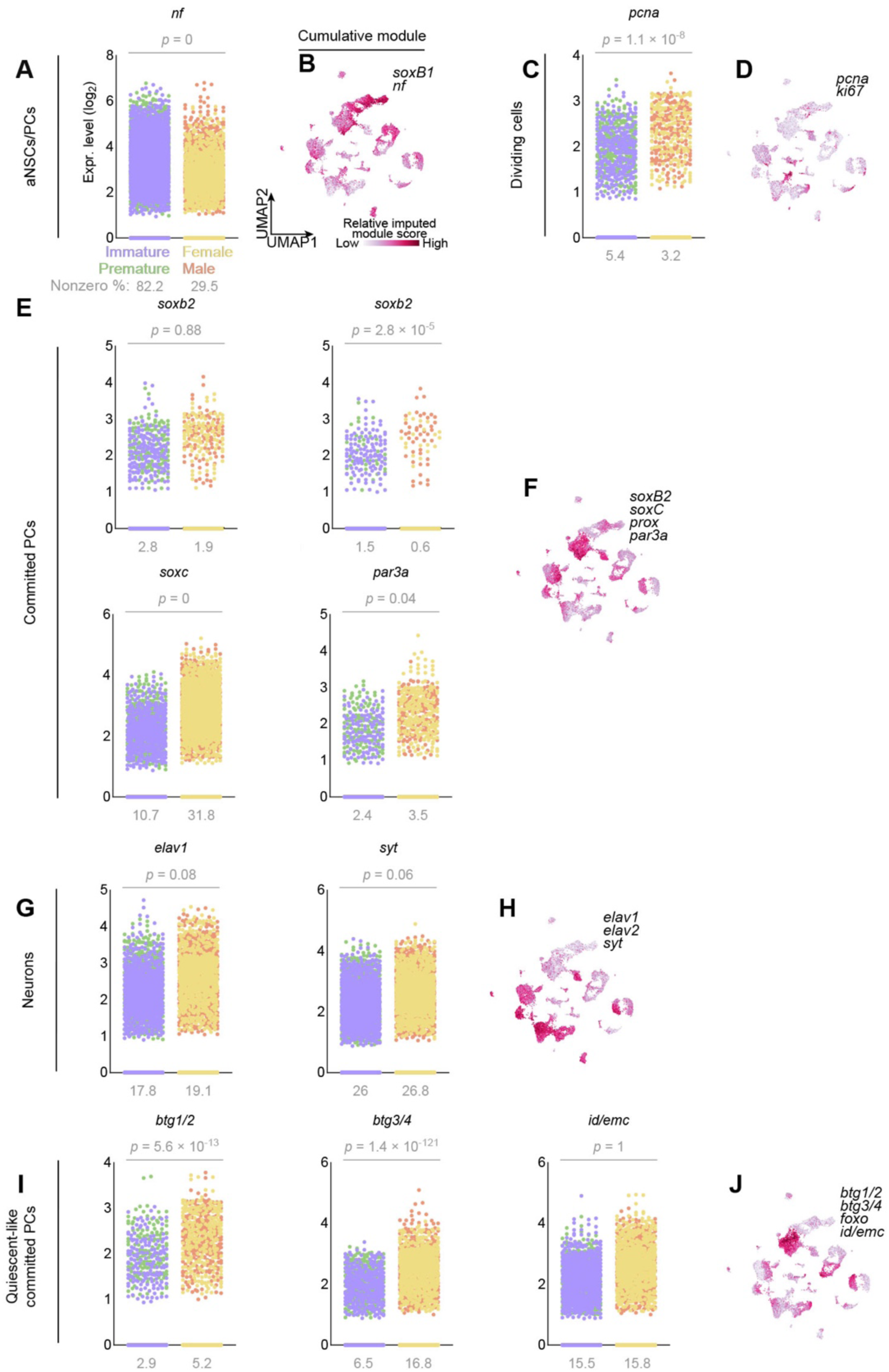
Expression of genes characteristic for distinct stages of the neurogenic cascade. Scatter plots and cumulative gene modules showing respective individual and cumulative expression of genes associated with (**A-B**) aNSCs/PCs, (**C-D**) proliferation, (**E-F**) neural commitment, (**G-H**) post-mitotic neurons and (**I-J**) quiescence. Statistical significance of expression means of non-reproductive and reproductive libraries was calculated using a Wilcoxon Rank Sum test.

**Fig. S12.**
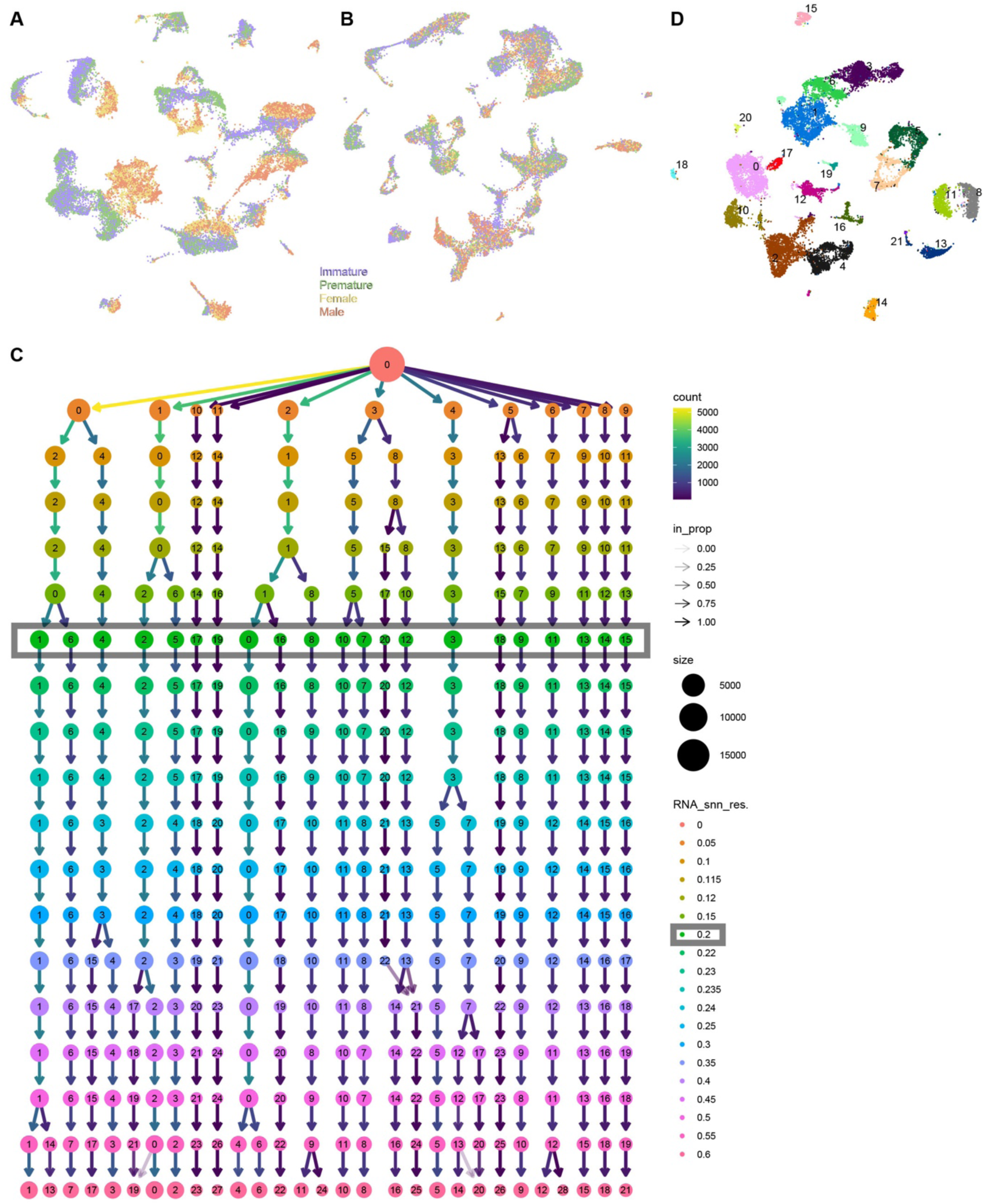
Integration and clustering quality controls of the brain atlas. Immature, premature, female and male libraries in their original size integrated following the (**A**) merge-integration and (**B**) standard integration pipeline, omitting downsampling. (**C**) Clustering tree representing increasing clustering resolutions (RNA_snn_res) and the resulting cluster numbers. (**D**) Cluster identities generated using the outlined clustering resolution in (**C**); the clusters correspond to cells colored with different shades of the same color within one overarching cluster identity in **Fig. 1D**.

**Fig. S13.**
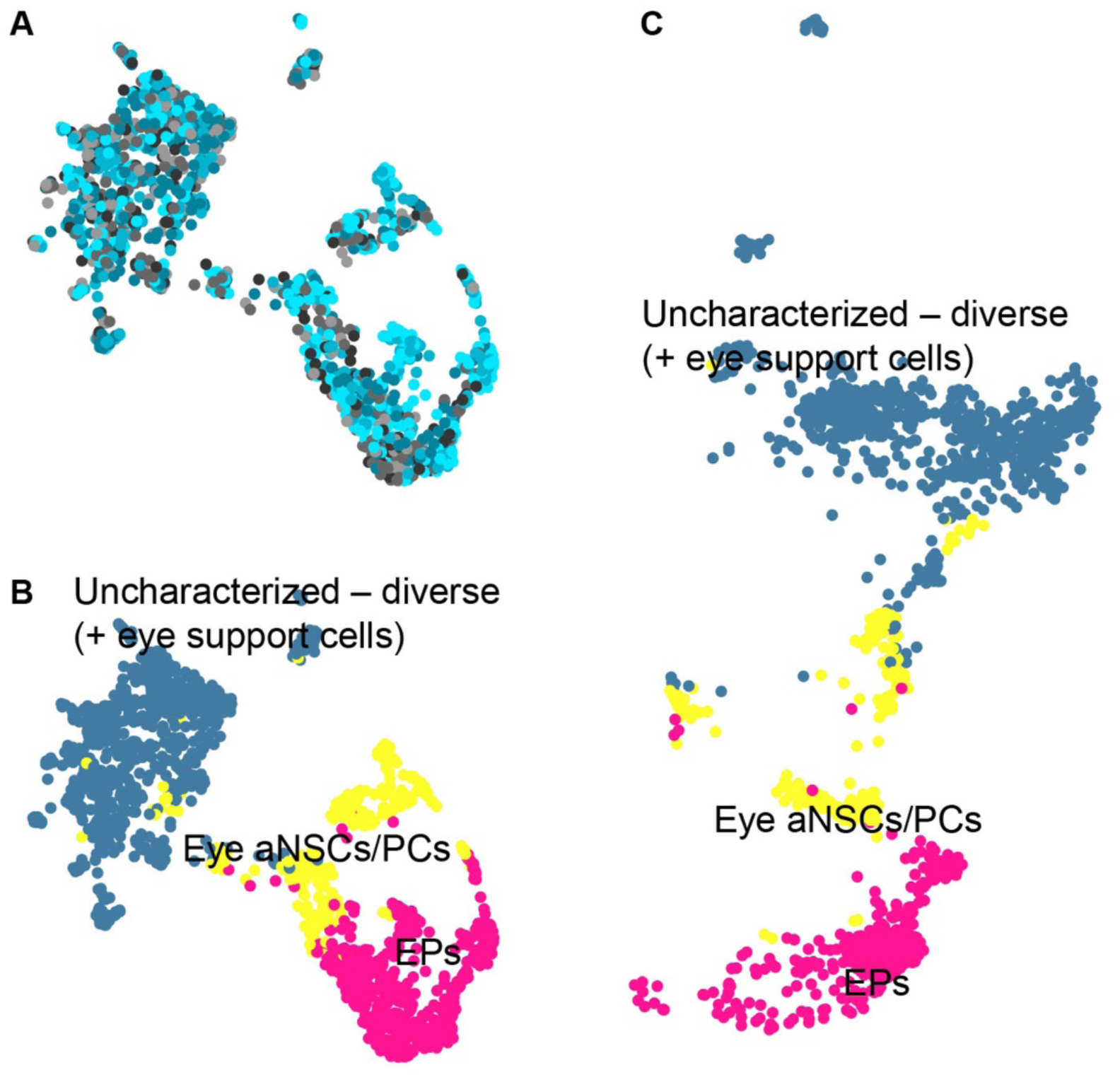
Predicted reduction and annotation of the eye subset of the *c-ops1^Δ8/Δ8^* and WT dataset. (**A**) Standard UMAP-reduction of the *c-ops1^Δ8/Δ8^* and WT eye- and transcriptomically related cell subset (query). Predicted overarching cluster identities on the (**B**) query and (**C**) predicted UMAP-reduction of the eye- and transcriptomically related cell subset, generated using the UMAP-coordinates and overarching cluster identities of the corresponding subset of the brain atlas as a reference, respectively.

**Table S1.** Synopsis of investigated *Platynereis* genes with predicted functions in neurogenesis or quiescence.

| *Platynereis* name | Human ortholog(s) | Genbank ID/ Source |
| --- | --- | --- |
| **Neurogenic and proneural genes** |  |  |
| *soxB1* | *sox1/sox2/sox3* | ANS60443.1 |
| *nf* | *gfap* | This study |
| *soxC* | *sox4/sox11/sox12* | ref. (2) |
| *prox* | *prox1* | ref. (2) |
| *par3a* | *pard3* | ref. (3) |
| *soxB2* | *sox14/sox21* | refs. (2, 4) |
| **Dividing cells** |  |  |
| *pcna* | *pcna* | HF935038.1 |
| *ki67* | *ki67* | This study |
| **Postmitotic Neurons** |  |  |
| *elav1* | *elavl1* | ref. (5) |
| *elav2* | *elavl2* | This study |
| *syt* | *syt* | ref. (6) |
| **Quiescent-like cell markers** |  |  |
| *btg1/2* | *btg1/btg2* | This study |
| *btg3/4* | *btg3/btg4* | This study |
| *foxo* | *foxo1/foxo3/foxo4/foxo6* | This study |

## References

1. H. Grandel, M. Brand, Comparative aspects of adult neural stem cell activity in vertebrates. Dev. Genes Evol. 223, 131–147 (2013).

2. F. Nottebohm, M. E. Nottebohm, L. Crane, Developmental and seasonal changes in canary song and their relation to changes in the anatomy of song-control nuclei. Behav. Neural Biol. 46, 445–471 (1986).

3. S. A. Goldman, F. Nottebohm, Neuronal production, migration, and differentiation in a vocal control nucleus of the adult female canary brain. Proc. Natl. Acad. Sci. 80, 2390–2394 (1983).

4. W. A. Harris, M. Perron, Molecular recapitulation: the growth of the vertebrate retina. Int. J. Dev. Biol. 42, 299–304 (1998).

5. E. Tsingos, B. Höckendorf, T. Sütterlin, S. Kirchmaier, N. Grabe, L. Centanin, J. Wittbrodt, Retinal stem cells modulate proliferative parameters to coordinate post-embryonic morphogenesis in the eye of fish. Elife 8, e42646 (2019).

6. A. Miles, V. Tropepe, Retinal Stem Cell ‘Retirement Plans’: Growth, Regulation and Species Adaptations in the Retinal Ciliary Marginal Zone. Int J Mol Sci 22, 6528 (2021).

7. I. Hernández-Núñez, A. Quelle-Regaldie, L. Sánchez, F. Adrio, E. Candal, A. Barreiro-Iglesias, Decline in Constitutive Proliferative Activity in the Zebrafish Retina with Ageing. Int. J. Mol. Sci. 22, 11715 (2021).

8. N. Sokolova, L. Zilova, J. Wittbrodt, Unravelling the link between embryogenesis and adult stem cell potential in the ciliary marginal zone: A comparative study between mammals and teleost fish. Cells Dev. 174, 203848 (2023).

9. I. Hernández-Núñez, D. Robledo, H. Mayeur, S. Mazan, L. Sánchez, F. Adrio, A. Barreiro-Iglesias, E. Candal, Loss of Active Neurogenesis in the Adult Shark Retina. Front. Cell Dev. Biol. 9, 628721 (2021).

10. G. Ming, H. Song, Adult Neurogenesis in the Mammalian Brain: Significant Answers and Significant Questions. Neuron 70, 687–702 (2011).

11. N. Urbán, I. M. Blomfield, F. Guillemot, Quiescence of Adult Mammalian Neural Stem Cells: A Highly Regulated Rest. Neuron 104, 834–848 (2019).

12. B. Trebels, S. Dippel, M. Schaaf, K. Balakrishnan, E. A. Wimmer, J. Schachtner, Adult neurogenesis in the mushroom bodies of red flour beetles (Tribolium castaneum, Herbst) is influenced by the olfactory environment. Sci. Rep. 10, 1090 (2020).

13. A. Hansen, M. Schmidt, Influence of season and environment on adult neurogenesis in the central olfactory pathway of the shore crab, Carcinus maenas. Brain Res. 1025, 85–97 (2004).

14. C. Bertapelle, G. Polese, A. D. Cosmo, Enriched Environment Increases PCNA and PARP1 Levels in Octopus vulgaris Central Nervous System: First Evidence of Adult Neurogenesis in Lophotrochozoa. J. Exp. Zoöl. Part B: Mol. Dev. Evol. 328, 347–359 (2017).

15. C. A. Penick, M. Ghaninia, K. L. Haight, C. Opachaloemphan, H. Yan, D. Reinberg, J. Liebig, Reversible plasticity in brain size, behaviour and physiology characterizes caste transitions in a socially flexible ant (Harpegnathos saltator). Proc. R. Soc. B 288, 20210141 (2021).

16. J. Gospocic, K. M. Glastad, L. Sheng, E. J. Shields, S. L. Berger, R. Bonasio, Kr-h1 maintains distinct caste-specific neurotranscriptomes in response to socially regulated hormones. Cell 184, 5807–5823.e14 (2021).

17. D. Arendt, K. Tessmar-Raible, H. Snyman, A. W. Dorresteijn, J. Wittbrodt, Ciliary Photoreceptors with a Vertebrate-Type Opsin in an Invertebrate Brain. Science 306, 869–871 (2004).

18. M. Zurl, B. Poehn, D. Rieger, S. Krishnan, D. Rokvic, V. B. V. Rajan, E. Gerrard, M. Schlichting, L. Orel, A. Ćorić, R. J. Lucas, E. Wolf, C. Helfrich-Förster, F. Raible, K. Tessmar-Raible, Two light sensors decode moonlight versus sunlight to adjust a plastic circadian/circalunidian clock to moon phase. Proc National Acad Sci 119, e2115725119 (2022).

19. H. Tsukamoto, I.-S. Chen, Y. Kubo, Y. Furutani, A ciliary opsin in the brain of a marine annelid zooplankton is ultraviolet-sensitive, and the sensitivity is tuned by a single amino acid residue. J. Biol. Chem. 292, 12971–12980 (2017).

20. V. B. V. Rajan, N. S. Häfker, E. Arboleda, B. Poehn, T. Gossenreiter, E. Gerrard, M. Hofbauer, C. Mühlestein, A. Bileck, C. Gerner, M. R. d’Alcala, M. C. Buia, M. Hartl, R. J. Lucas, K. Tessmar-Raible, Seasonal variation in UVA light drives hormonal and behavioural changes in a marine annelid via a ciliary opsin. Nat Ecol Evol 5, 204–218 (2021).

21. C. Verasztó, M. Gühmann, H. Jia, V. B. V. Rajan, L. A. Bezares-Calderón, C. Piñeiro-Lopez, N. Randel, R. Shahidi, N. K. Michiels, S. Yokoyama, K. Tessmar-Raible, G. Jékely, Ciliary and rhabdomeric photoreceptor-cell circuits form a spectral depth gauge in marine zooplankton. Elife 7, e36440 (2018).

22. B. J. Eriksson, D. Fredman, G. Steiner, A. Schmid, Characterisation and localisation of the opsin protein repertoire in the brain and retinas of a spider and an onychophoran. BMC Evol. Biol. 13, 186 (2013).

23. O. Vöcking, A. Macias-Muñoz, S. J. Jaeger, T. H. Oakley, Deep Diversity: Extensive Variation in the Components of Complex Visual Systems across Animals. Cells 11, 3966 (2022).

24. C. Hauenschild, Abhängigkeit der Regenerationsleistung von der inneren Sekretion im Prostomium bei Platynereis dumerilii. Z. für Naturforsch. B 15, 52–55 (1960).

25. C. Hauenschild, Der hormonale Einfluss des Gehirns auf die sexuelle Entwicklung bei dem polychaeten Platynereis dumerilii. Gen Comp Endocr 6, 26–73 (1966).

26. C. Hauenschild, A. Fischer, “Platynereis dumerilii. Mikroskopische Anatomie, Fortpflanzung und Entwicklung [Platynereis dumerilii. Microscopical anatomy, reproduction and development]” in Grosses Zoologisches Praktikum (1969).

27. D. K. Hofmann, Regeneration and endocrinology in the polychaete Platynereis dumerilii. An experimental and structural study. *Wilhelm Roux’* Archiv für Entwicklungsmechanik der Organismen 180, 47–71 (1976).

28. S. Schenk, S. C. Bannister, F. J. Sedlazeck, D. Anrather, B. Q. Minh, A. Bileck, M. Hartl, A. von Haeseler, C. Gerner, F. Raible, K. Tessmar-Raible, Combined transcriptome and proteome profiling reveals specific molecular brain signatures for sex, maturation and circalunar clock phase. Elife 8, e41556 (2019).

29. L. Culig, X. Chu, V. A. Bohr, Neurogenesis in aging and age-related neurodegenerative diseases. Ageing Res. Rev. 78, 101636 (2022).

30. M. Iglesias, D. A. Felix, Ó. Gutiérrez-Gutiérrez, M. del M. D. Miguel-Bonet, S. Sahu, B. Fernández-Varas, R. Perona, A. A. Aboobaker, I. Flores, C. González-Estévez, Downregulation of mTOR Signaling Increases Stem Cell Population Telomere Length during Starvation of Immortal Planarians. Stem Cell Rep. 13, 405–418 (2019).

31. Á. Varley, H. R. Horkan, E. T. McMahon, G. Krasovec, U. Frank, Pluripotent, germ cell competent adult stem cells underlie cnidarian regenerative ability and clonal growth. Curr. Biol. 33, 1883–1892.e3 (2023).

32. J. Steger, A. G. Cole, A. Denner, T. Lebedeva, G. Genikhovich, A. Ries, R. Reischl, E. Taudes, M. Lassnig, U. Technau, Single-cell transcriptomics identifies conserved regulators of neuroglandular lineages. Cell Reports 40, 111370 (2022).

33. Y. Hao, S. Hao, E. Andersen-Nissen, W. M. Mauck, S. Zheng, A. Butler, M. J. Lee, A. J. Wilk, C. Darby, M. Zager, P. Hoffman, M. Stoeckius, E. Papalexi, E. P. Mimitou, J. Jain, A. Srivastava, T. Stuart, L. M. Fleming, B. Yeung, A. J. Rogers, J. M. McElrath, C. A. Blish, R. Gottardo, P. Smibert, R. Satija, Integrated analysis of multimodal single-cell data. Cell 184, 3573–3587.e29 (2021).

34. P. Kerner, E. Simionato, M. L. Gouar, M. Vervoort, Orthologs of key vertebrate neural genes are expressed during neurogenesis in the annelid Platynereis dumerilii. Evol Dev 11, 513–524 (2009).

35. A. W. Stockinger, L. Adelmann, M. Fahrenberger, C. Ruta, D. Özpolat, N. Milivojev, G. Balavoine, F. Raible, Molecular profile, source and lineage restriction of stem cells in an annelid regeneration model. Nat Comms, under revision.

36. K. Stamatiou, P. Vagnarelli, Chromosome clustering in mitosis by the nuclear protein Ki-67. Biochem. Soc. Trans. 49, 2767–2776 (2021).

37. A. Fischer, J. Brökelmann, [The eye of Platynereis dumerilii (Polychaeta): its fine structure in ontogenetic and adaptive change]. Zeitschrift Für Zellforschung Und Mikroskopische Anatomie Vienna Austria 1948 71, 217–44 (1966).

38. R. Revilla-i-Domingo, V. B. V. Rajan, M. Waldherr, G. Prohaczka, H. Musset, L. Orel, E. Gerrard, M. Smolka, A. Stockinger, M. Farlik, R. J. Lucas, F. Raible, K. Tessmar-Raible, Characterization of cephalic and non-cephalic sensory cell types provides insight into joint photo- and mechanoreceptor evolution. Elife 10, e66144 (2021).

39. B. Rhode, Development and differentiation of the eye in Platynereis dumerilii (Annelida, Polychaeta). J Morphol 212, 71–85 (1992).

40. R. Mollicone, S. E. H. Moore, N. Bovin, M. Garcia-Rosasco, J.-J. Candelier, I. Martinez-Duncker, R. Oriol, Activity, Splice Variants, Conserved Peptide Motifs, and Phylogeny of Two New α1,3-Fucosyltransferase Families (FUT10 and FUT11)*. J. Biol. Chem. 284, 4723–4738 (2009).

41. T. Ayers, H. Tsukamoto, M. Gühmann, V. B. V. Rajan, K. Tessmar-Raible, A Go-type opsin mediates the shadow reflex in the annelid Platynereis dumerilii. Bmc Biol 16, 41 (2018).

42. M. A. Tosches, D. Bucher, P. Vopalensky, D. Arendt, Melatonin Signaling Controls Circadian Swimming Behavior in Marine Zooplankton. Cell 159, 46–57 (2014).

43. J. Cao, M. Spielmann, X. Qiu, X. Huang, D. M. Ibrahim, A. J. Hill, F. Zhang, S. Mundlos, L. Christiansen, F. J. Steemers, C. Trapnell, J. Shendure, The single-cell transcriptional landscape of mammalian organogenesis. Nature 566, 496–502 (2019).

44. L. Centanin, J.-J. Ander, B. Hoeckendorf, K. Lust, T. Kellner, I. Kraemer, C. Urbany, E. Hasel, W. A. Harris, B. D. Simons, J. Wittbrodt, Exclusive multipotency and preferential asymmetric divisions in post-embryonic neural stem cells of the fish retina. *Dev. (Camb.*, Engl.) 141, 3472–3482 (2014).

45. Y. Wan, A. D. Almeida, S. Rulands, N. Chalour, L. Muresan, Y. Wu, B. D. Simons, J. He, W. A. Harris, The ciliary marginal zone of the zebrafish retina: clonal and time-lapse analysis of a continuously growing tissue. Development 143, 1099–1107 (2016).

46. D. Arendt, K. Tessmar, M.-I. M. de Campos-Baptista, A. Dorresteijn, J. Wittbrodt, Development of pigment-cup eyes in the polychaete Platynereis dumerilii and evolutionary conservation of larval eyes in Bilateria. Dev Camb Engl 129, 1143–54 (2002).

47. B. Backfisch, V. B. V. Rajan, R. M. Fischer, C. Lohs, E. Arboleda, K. Tessmar-Raible, F. Raible, Stable transgenesis in the marine annelid Platynereis dumerilii sheds new light on photoreceptor evolution. Proc National Acad Sci 110, 193–198 (2013).

48. N. Randel, L. A. Bezares-Calderón, M. Gühmann, R. Shahidi, G. Jékely, Expression Dynamics and Protein Localization of Rhabdomeric Opsins in Platynereis Larvae. Integr Comp Biol 53, 7–16 (2013).

49. P. Codega, V. Silva-Vargas, A. Paul, A. R. Maldonado-Soto, A. M. DeLeo, E. Pastrana, F. Doetsch, Prospective Identification and Purification of Quiescent Adult Neural Stem Cells from Their In Vivo Niche. Neuron 82, 545–559 (2014).

50. N. Urbán, Could a Different View of Quiescence Help Us Understand How Neurogenesis Is Regulated? Front. Neurosci. 16, 878875 (2022).

51. A. B. Nakama, H.-C. Chou, S. Q. Schneider, The asymmetric cell division machinery in the spiral-cleaving egg and embryo of the marine annelid Platynereis dumerilii. BMC Dev. Biol. 17, 16 (2017).

52. J. Paik, Z. Ding, R. Narurkar, S. Ramkissoon, F. Muller, W. S. Kamoun, S.-S. Chae, H. Zheng, H. Ying, J. Mahoney, D. Hiller, S. Jiang, A. Protopopov, W. H. Wong, L. Chin, K. L. Ligon, R. A. DePinho, FoxOs Cooperatively Regulate Diverse Pathways Governing Neural Stem Cell Homeostasis. Cell Stem Cell 5, 540–553 (2009).

53. J. Andrade, C. Shi, A. S. H. Costa, J. Choi, J. Kim, A. Doddaballapur, T. Sugino, Y. T. Ong, M. Castro, B. Zimmermann, M. Kaulich, S. Guenther, K. Wilhelm, Y. Kubota, T. Braun, G. Y. Koh, A. R. Grosso, C. Frezza, M. Potente, Control of endothelial quiescence by FOXO-regulated metabolites. Nat. Cell Biol. 23, 413–423 (2021).

54. S. S. Hwang, J. Lim, Z. Yu, P. Kong, E. Sefik, H. Xu, C. C. D. Harman, L. K. Kim, G. R. Lee, H.-B. Li, R. A. Flavell, mRNA destabilization by BTG1 and BTG2 maintains T cell quiescence. Science 367, 1255–1260 (2020).

55. L. Wei, E. C. Lai, Regulation of the Alternative Neural Transcriptome by ELAV/Hu RNA Binding Proteins. Front. Genet. 13, 848626 (2022).

56. M. R. Bowers, N. E. Reist, Synaptotagmin: Mechanisms of an electrostatic switch. Neurosci. Lett. 722, 134834 (2020).

57. A. Ćorić, A. W. Stockinger, P. Schaffer, D. Rokvić, K. Tessmar-Raible, F. Raible, A Fast And Versatile Method for Simultaneous HCR, Immunohistochemistry And Edu Labeling (SHInE). Integr Comp Biol, doi: 10.1093/icb/icad007 (2023).

58. C. T. Miller, M. E. Hale, H. Okano, S. Okabe, P. Mitra, Comparative Principles for Next-Generation Neuroscience. Front. Behav. Neurosci. 13, 12 (2019).

59. K. Tessmar-Raible, F. Raible, F. Christodoulou, K. Guy, M. Rembold, H. Hausen, D. Arendt, Conserved Sensory-Neurosecretory Cell Types in Annelid and Fish Forebrain: Insights into Hypothalamus Evolution. Cell 129, 1389–1400 (2007).

60. D. Arendt, P. Y. Bertucci, K. Achim, J. M. Musser, Evolution of neuronal types and families. Curr Opin Neurobiol 56, 144–152 (2019).

61. D. Arendt, M. A. Tosches, H. Marlow, From nerve net to nerve ring, nerve cord and brain — evolution of the nervous system. Nat Rev Neurosci 17, 61–72 (2016).

62. A. S. Denes, G. Jékely, P. R. H. Steinmetz, F. Raible, H. Snyman, B. Prud’homme, D. E. K. Ferrier, G. Balavoine, D. Arendt, Molecular Architecture of Annelid Nerve Cord Supports Common Origin of Nervous System Centralization in Bilateria. Cell 129, 277–288 (2007).

63. Y. Liang, A. M. Carrillo-Baltodano, J. M. Martín-Durán, Emerging trends in the study of spiralian larvae. Evol. Dev., e12459 (2023).

64. B. D. Özpolat, N. Randel, E. A. Williams, L. A. Bezares-Calderón, G. Andreatta, G. Balavoine, P. Y. Bertucci, D. E. K. Ferrier, M. C. Gambi, E. Gazave, M. Handberg-Thorsager, J. Hardege, C. Hird, Y.-W. Hsieh, J. Hui, K. N. Mutemi, S. Q. Schneider, O. Simakov, H. M. Vergara, M. Vervoort, G. Jékely, K. Tessmar-Raible, F. Raible, D. Arendt, The Nereid on the rise: Platynereis as a model system. Evodevo 12, 10 (2021).

65. F. Tirone, S. Farioli-Vecchioli, L. Micheli, M. Ceccarelli, L. Leonardi, Genetic control of adult neurogenesis: interplay of differentiation, proliferation and survival modulates new neurons function, and memory circuits. Front. Cell. Neurosci. 7, 59 (2013).

66. J. D. Norton, ID helix-loop-helix proteins in cell growth, differentiation and tumorigenesis. J. Cell Sci. 113, 3897–3905 (2000).

67. K. Li, N. E. Baker, Regulation of the Drosophila ID protein Extra macrochaetae by proneural dimerization partners. eLife 7, e33967 (2018).

68. D. W. Golding, E. Yuwono, Latent capacities for gametogenic cycling in the semelparous invertebrate Nereis. Proc National Acad Sci 91, 11777–11781 (1994).

69. A. Fischer, “Reproductive Strategies and Developmental Patterns in Annelids” in Reproductive Strategies and Developmental Patterns in Annelids (Reproductive Strategies and Developmental Patterns in Annelids, 1999; http://link.springer.com/10.1007/978-94-017-2887-4_1) *Reproductive Strategies and Developmental Patterns in Annelids*, pp. 1–20.

70. P. H. O’Farrell, Quiescence: early evolutionary origins and universality do not imply uniformity. Philos. Trans. R. Soc. B: Biol. Sci. 366, 3498–3507 (2011).

71. F. R. Napoli, C. M. Daly, S. Neal, K. J. McCulloch, A. R. Zaloga, A. Liu, K. M. Koenig, Cephalopod retinal development shows vertebrate-like mechanisms of neurogenesis. Curr. Biol. 32, 5045–5056.e3 (2022).

72. M. D. Ramirez, A. N. Pairett, M. S. Pankey, J. M. Serb, D. I. Speiser, A. J. Swafford, T. H. Oakley, The Last Common Ancestor of Most Bilaterian Animals Possessed at Least Nine Opsins. Genome Biol Evol 8, 3640–3652 (2016).

73. A. Mat, H. H. Vu, E. Wolf, K. Tessmar-Raible, All Light, Everywhere? Photoreceptors at Nonconventional Sites. Physiology 39, 30–43 (2024).

74. B. Poehn, S. Krishnan, M. Zurl, A. Coric, D. Rokvic, N. S. Häfker, E. Jaenicke, E. Arboleda, L. Orel, F. Raible, E. Wolf, K. Tessmar-Raible, A Cryptochrome adopts distinct moon- and sunlight states and functions as sun-versus moonlight interpreter in monthly oscillator entrainment. Nat Commun 13, 5220 (2022).

75. H. Tsukamoto, Y. Kubo, A self-inactivating invertebrate opsin optically drives biased signaling toward Gβγ-dependent ion channel modulation. Proc. Natl. Acad. Sci. United States Am. 120, e2301269120 (2023).

76. H. García-Castro, N. J. Kenny, M. Iglesias, P. Álvarez-Campos, V. Mason, A. Elek, A. Schönauer, V. A. Sleight, J. Neiro, A. Aboobaker, J. Permanyer, M. Irimia, A. Sebé-Pedrós, J. Solana, ACME dissociation: a versatile cell fixation-dissociation method for single-cell transcriptomics. Genome Biol. 22, 89 (2021).

77. K. N. Mutemi, O. Simakov, L. Pan, L. Santangeli, R. Null, M. Handberg-Thorsager, B. C. Vellutini, T. Larsson, E. Savage, M. O. Lopez, R. Hercog, J. Provaznik, D. Ordoñez-Rueda, N. Azevedo, E. Gazave, M. Vervoort, P. Tomancak, W. Tan, S. Winkler, V. Benes, J. Hui, C. Helm, B. D. Özpolat, D. Arendt, A genome resource for the marine annelid Platynereis dumerilii. bioRxiv, 2024.06.21.600153 (2024).

78. E. Becht, L. McInnes, J. Healy, C.-A. Dutertre, I. W. H. Kwok, L. G. Ng, F. Ginhoux, E. W. Newell, Dimensionality reduction for visualizing single-cell data using UMAP. Nat. Biotechnol. 37, 38–44 (2019).

79. L. Zappia, A. Oshlack, Clustering trees: a visualization for evaluating clusterings at multiple resolutions. GigaScience 7, giy083 (2018).

80. Stockinger, L. Adelmann, M. Fahrenmberger, C. Ruta, B. D. Özpolat, N. Milivojev, G. Balavoine, F. Raible, Molecular profile, source and lineage restriction of stem cells in an annelid regeneration model. Nat Comms.

81. T. Lubiana, H. Nakaya, Using coexpression to explore cell-type diversity with the fcoex package. bioRxiv, 2021.12.07.471603 (2021).

82. G. C. Linderman, J. Zhao, M. Roulis, P. Bielecki, R. A. Flavell, B. Nadler, Y. Kluger, Zero-preserving imputation of single-cell RNA-seq data. Nat. Commun. 13, 192 (2022).

83. J. Schindelin, I. Arganda-Carreras, E. Frise, V. Kaynig, M. Longair, T. Pietzsch, S. Preibisch, C. Rueden, S. Saalfeld, B. Schmid, J.-Y. Tinevez, D. J. White, V. Hartenstein, K. Eliceiri, P. Tomancak, A. Cardona, Fiji: an open-source platform for biological-image analysis. Nat. Methods 9, 676–682 (2012).

84. E. Kuehn, D. S. Clausen, R. W. Null, B. M. Metzger, A. D. Willis, B. D. Özpolat, Segment number threshold determines juvenile onset of germline cluster expansion in Platynereis dumerilii. J Exp Zoology Part B Mol Dev Evol 338, 225–240 (2022).

85. M. Čapek, J. Janáček, L. Kubínová, Methods for compensation of the light attenuation with depth of images captured by a confocal microscope. Microsc. Res. Tech. 69, 624–635 (2006).

86. S. X. Ge, D. Jung, R. Yao, ShinyGO: a graphical gene-set enrichment tool for animals and plants. Bioinformatics 36, 2628–2629 (2019).

87. S. Carbon, A. Ireland, C. J. Mungall, S. Shu, B. Marshall, S. Lewis, the A. Hub, the W. P. W. Group, AmiGO: online access to ontology and annotation data. Bioinformatics 25, 288–289 (2008).

88. N. S. Häfker, L. Holcik, A. M. Mat, A. Ćorić, K. Vadiwala, I. Beets, A. W. Stockinger, C. E. Atria, S. Hammer, R. Revilla-i-Domingo, L. Schoofs, F. Raible, K. Tessmar-Raible, Molecular circadian rhythms are robust in marine annelids lacking rhythmic behavior. PLOS Biol. 22, e3002572 (2024).

89. L.-T. Nguyen, H. A. Schmidt, A. von Haeseler, B. Q. Minh, IQ-TREE: A Fast and Effective Stochastic Algorithm for Estimating Maximum-Likelihood Phylogenies. Mol Biol Evol 32, 268– 274 (2015).

90. S. Kalyaanamoorthy, B. Q. Minh, T. K. F. Wong, A. von Haeseler, L. S. Jermiin, ModelFinder: fast model selection for accurate phylogenetic estimates. Nat Methods 14, 587– 589 (2017).

91. D. T. Hoang, O. Chernomor, A. von Haeseler, B. Q. Minh, L. S. Vinh, UFBoot2: Improving the Ultrafast Bootstrap Approximation. Mol. Biol. Evol. 35, 518–522 (2018).

92. I. Letunic, P. Bork, Interactive Tree Of Life (iTOL) v5: an online tool for phylogenetic tree display and annotation. Nucleic Acids Res. 49, W293–W296 (2021).

93. I. Letunic, S. Khedkar, P. Bork, SMART: recent updates, new developments and status in 2020. Nucleic Acids Res. 49, D458–D460 (2020).

## Supplementary references

1. R. Revilla-i-Domingo, V. B. V. Rajan, M. Waldherr, G. Prohaczka, H. Musset, L. Orel, E. Gerrard, M. Smolka, A. Stockinger, M. Farlik, R. J. Lucas, F. Raible, K. Tessmar-Raible, Characterization of cephalic and non-cephalic sensory cell types provides insight into joint photo- and mechanoreceptor evolution. Elife 10, e66144 (2021).

2. P. Kerner, E. Simionato, M. L. Gouar, M. Vervoort, Orthologs of key vertebrate neural genes are expressed during neurogenesis in the annelid Platynereis dumerilii. Evol Dev 11, 513–524 (2009).

3. A. B. Nakama, H.-C. Chou, S. Q. Schneider, The asymmetric cell division machinery in the spiral-cleaving egg and embryo of the marine annelid Platynereis dumerilii. BMC Dev. Biol. 17, 16 (2017).

4. Stockinger, L. Adelmann, M. Fahrenmberger, C. Ruta, B. D. Özpolat, N. Milivojev, G. Balavoine, F. Raible, Molecular profile, source and lineage restriction of stem cells in an annelid regeneration model. Nat Comms.

5 A. S. Denes, G. Jékely, P. R. H. Steinmetz, F. Raible, H. Snyman, B. Prud’homme, D. E. K. Ferrier, G. Balavoine, D. Arendt, Molecular Architecture of Annelid Nerve Cord Supports Common Origin of Nervous System Centralization in Bilateria. Cell 129, 277–288 (2007).

6. K. Tessmar-Raible, F. Raible, F. Christodoulou, K. Guy, M. Rembold, H. Hausen, D. Arendt, Conserved Sensory-Neurosecretory Cell Types in Annelid and Fish Forebrain: Insights into Hypothalamus Evolution. Cell 129, 1389–1400 (2007).

